# Epithelial antigen presentation controls commensal-specific intraepithelial T-cells in the gut

**DOI:** 10.1101/2022.08.21.504672

**Authors:** Tomas Brabec, Martin Schwarzer, Katarína Kováčová, Martina Dobešová, Dagmar Schierová, Jiří Březina, Iva Pacáková, Dagmar Šrůtková, Osher Ben-Nun, Yael Goldfarb, Iva Šplíchalová, Michal Kolář, Jakub Abramson, Dominik Filipp, Jan Dobeš

## Abstract

The expression of MHCII by intestinal epithelial cells (IEC) determines the severity of intestinal immunopathological reactions. However, the function of MHCII on IEC under homeostatic conditions remains elusive. Here we report that MHCII expression on IECs is a hallmark of an adaptive wave of homeostatic intestinal immune responses to commensal segmented filamentous bacteria (SFB). Focusing on SFB-driven responses, we describe the expression pattern of MHCII and the associated antigen processing machinery among IEC subpopulations along with the cellular network that regulates MHCII induction. Furthermore, we show that SFB induce the accumulation of SFB-specific intraepithelial lymphocytes (IELs) that originate from conventional CD4^+^ T-cells. Importantly, induced IELs are dependent on the epithelial MHCII. Finally, we demonstrate that both epithelial MHCII and the IEL functionality regulate the epithelial turnover. This study describes the organization of a commensal-targeted, IEL-driven immune response that is controlled by IEC antigen presentation and ultimately regulates IEC turnover.

## Introduction

Intestinal epithelial cells (IECs) form a single-layered intestinal epithelium, which facilitates the absorption of nutrients and water and is critical for restraining the intestinal microbiota and pathogens in the gut lumen. Loss of intestinal epithelium barrier integrity leads to the translocation of luminal components to the host body and may result in the development of severe inflammatory intestinal disorders due to the break-down of intestinal homeostasis (Bevins and Salzman, 2011; Maloy and Powrie, 2011).

It has been recognized for a long time that IEC can express the major histocompatibility complex class II (MHCII) molecules and function as antigen presenting cells with the capacity to activate CD4^+^ T-cells (Arnaud-Battandier et al., 1986; Gorvel et al., 1984; Scott et al., 1980). More recently, MHCII^+^ IECs were suggested to function as non-classical antigen presenting cells, which can modulate the severity of induced colitis in mice (Jamwal et al., 2020), as well as of graft-versus-host disease (Koyama et al., 2019) or intestinal tumorigenesis (Beyaz et al., 2021). This suggests that the IEC-dependent presentation of antigens in the MHCII context may contribute to CD4^+^ T cell-mediated adaptive immune responses. Additional studies revealed that the MHCII expression on IECs is partially regulated by the circadian clock and dietary intake rhythmicity (Tuganbaev et al., 2020) as well as by diet composition (Beyaz *et al*., 2021). Moreover, the MHCII expression on intestinal stem cells was implicated in organizing the stem cell differentiation potential to distinct epithelial cell lineages (Biton et al., 2018). However, the impact of MHCII^+^ IECs on T-cell response as well as the mechanism that conveys the function of MHCII^+^ IECs under steady-state conditions remains unclear since most of the previous studies focused on physiologically extreme situations.

The majority of luminal microbiota is prevented from direct adhesion to the apical side of IECs by the mucus layer, which also contains a gradient of IgA antibodies of different specificities and various antimicrobial peptides. In contrast, segmented filamentous bacteria (SFB), as well as other microbes, have developed a different life strategy, which involves an intimate interaction with IECs (Blumershine and Savage, 1977; Farkas et al., 2015; Klaasen et al., 1992). SFB are anaerobic, spore-forming, clostridia-like microorganisms that are under homeostatic conditions kept in check by immune surveillance mechanisms, preventing their penetration to the epithelial barrier and induction of a pathologic inflammatory response (Farkas *et al*., 2015; Talham et al., 1999). SFB uses a hook-like protrusion intercepted inside the IECs to facilitate their adhesion. This leads to a unique strategy of infection, where the SFB remains largely extracellular and only partially infects the inner intracellular space of the IECs without disrupting its plasma membrane. Interestingly, endocytic vesicles containing SFB cell wall-associated proteins are shed into the cytosol from the tip of the SFB-hook in the actin-dependent endocytic manner and shuttled through the endosomal-lysosomal compartment of IECs (Ladinsky et al., 2019) where MHCII-presentation machinery of the IECs is present. Tight adhesion of SFB to IEC induces a strong immune response in the intestine, which is characterized by the initiation of a T helper 17 (T_H_17) response (Atarashi et al., 2015; Gaboriau-Routhiau et al., 2009; Ivanov et al., 2009; Yang et al., 2014), including increased production of IL-17 and IL-22, as well as by increased production of IgA (Lecuyer et al., 2014). These responses in turn limit the overgrowth of SFB (Kumar et al., 2016; Shih et al., 2014; Suzuki et al., 2004).

The IECs are wired with the immune system through close physical and functional interactions mediated by several immune cell types located in the intraepithelial space of the gut, including a heterogeneous group of intraepithelial lymphocytes (IELs) (Cheroutre et al., 2011; Hayday et al., 2001). IELs are characterized by high expression of an adhesion integrin molecule CD103, which facilitates their interaction with E-cadherin on IECs (Cepek et al., 1994; Kilshaw and Murant, 1990). Most of IELs also express interaction partners of thymus leukemia antigen (TLA, encoded by the *H2-T3* gene), CD8αα homodimers (Leishman et al., 2001). IELs in general were implicated in immune protection against viruses (Masopust et al., 2006; Müller et al., 2000; Shires et al., 2001), bacteria (Mombaerts et al., 1993) or parasites (Chardès et al., 1994; Lepage et al., 1998). IELs can be broadly classified into two major sub-types – natural IELs and induced IELs (iIELs). While natural IELs are generated in the thymus, iIEL arise from activated, antigen-experienced conventional TCRαβ^+^ T-cells (Cheroutre *et al*., 2011). Although a portion of iIEL arise from CD4^+^ T-cell, their classical ThPOK-guarded CD4^+^ identity is shaken by the induction of Runx3-mediated suppression of this master regulator(Reis et al., 2013), leading to the loss of ThPOK expression (Mucida et al., 2013) and establishment of a potent CD8^+^ T-cell-like phenotype manifested by large cytolytic granules (Shires *et al*., 2001). Formerly CD4^+^ iIELs keep their MHCII-restriction although losing CD4^+^ T-cell identity (He et al., 2005; Mucida *et al*., 2013). During aging, exogenous stimuli induce a gradual accumulation of iIEL in the intestinal intraepithelial space (Umesaki et al., 1993). The homeostatic function of iIELs bearing MHCII-restricted αβTCR and cytotoxic granules remains rather poorly explored.

Here we report that MHCII-expression on IECs is strictly dependent on the age of the host and the colonization of the host’
ss gut by SFB. The SFB presence induces IFNγ production in T-cells, which in turn stimulates IECs to express MHCII. Moreover, SFB induce a switch of SFB-specific CD4^+^ T-cells to granzyme-expressing iIELs. The MHCII presence on IECs is essential for the proper positioning of SFB-specific iIELs in the gut intraepithelial space and regulation of the intestinal epithelium turnover under homeostatic conditions.

## Results

### Profiling of intestinal immune cells reveals different age-dependent kinetics of Th1, Th17 and cytotoxic T-cell responses in the intestine

Microbiota-derived cues fine-tune immune system responses and prevent the development of immunopathologies, such as allergy or inflammatory bowel disease (Gensollen et al., 2016; Kronman et al., 2012; Prioult and Nagler-Anderson, 2005). Recently, it was shown that these microbial cues must be delivered in mice around 3 weeks of age, in a specific time window during weaning. At this time point, mice start to consume solid food and this leads to profound changes in microbiota composition initiating a robust immune response (Al Nabhani et al., 2019). While it is known that MHCII expression on IECs affects the severity of various immunopathologies (Jamwal *et al*., 2020; Thelemann et al., 2014) and is dependent on intestinal microbiota (Van Der Kraak et al., 2021), very little is known about the role of MHCII^+^ IECs in regard to time-dependent microbiota responses. Specifically, it is still unclear when exactly IECs start to express MHCII, what specific signals induce it, or whether it conveys any of the time-dependent microbiota-induced stimuli. To better map the MHCII expression pattern and to elucidate potential transcriptional changes in IECs and gut-resident immune populations throughout aging, we performed single-cell RNA sequencing (scRNA-seq) of FACS sorted CD45^+^ immune cells residing in small intestinal (SI) lamina propria (LP), as well as FACS sorted CD45^−^ EpCAM^+^ SI-IEC derived from 3-, 4-, 5- and 6- weeks old wild-type (WT) littermate mice. Focusing first on SI-LP analysis, we performed clustering and sub-clustering analysis based on their transcription profile, defining intestinal immune cell types (Figures 1A-1C and S1).

**Figure 1.**
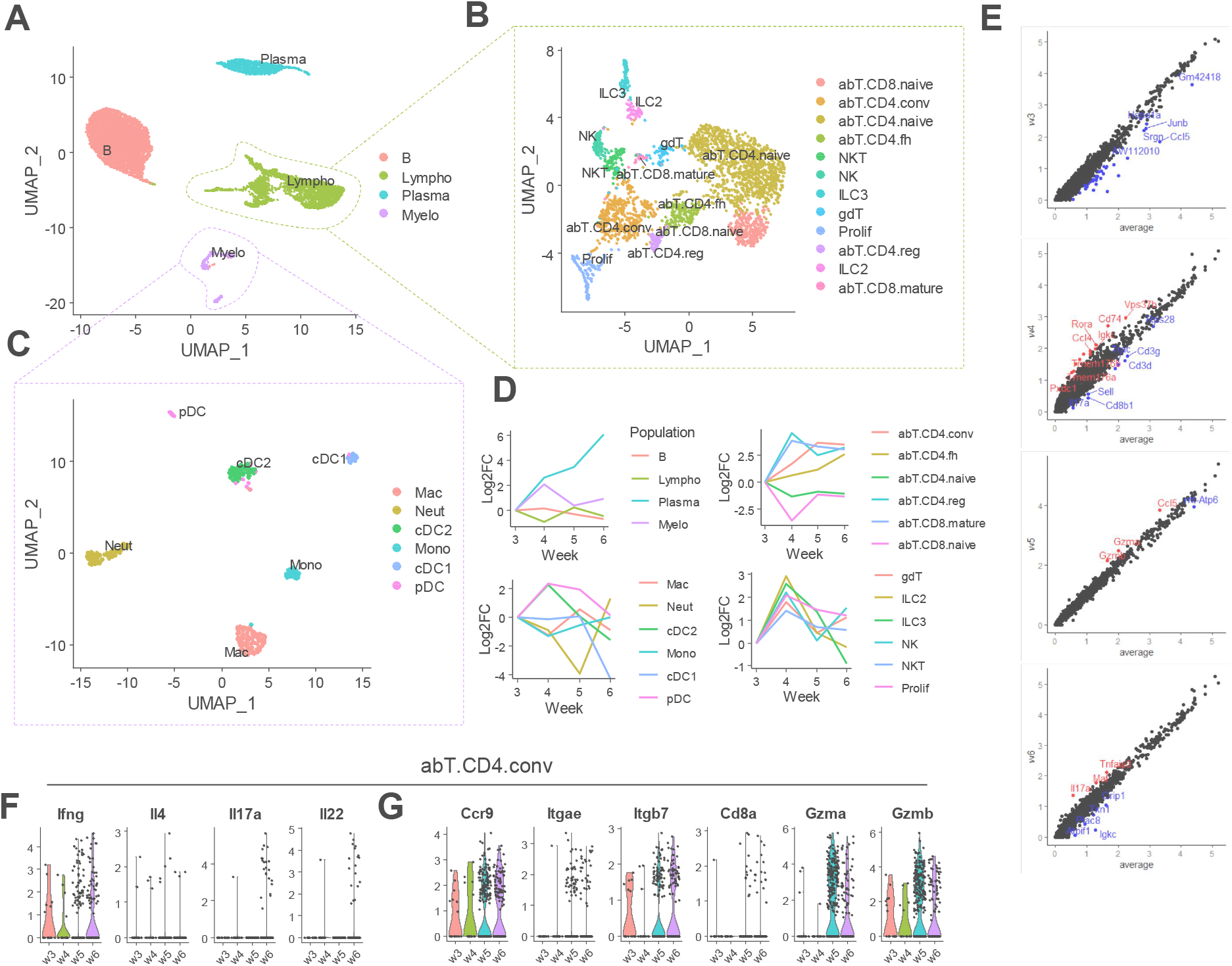
Profiling of lamina propria immune cells through mouse age. (A-G) scRNA-seq analysis of SI-LP hematopoietic cells from 3-, 4-, 5- and 6-weeks old WT littermate mice. (A) UMAP of SI-LP cells showing major clusters of B-cells (based on *Cd19, H2-Ab1* expression), plasma cells (*Sdc1, Igha*), other lymphoid cells (*Cd3d, Il7r*) and myeloid cells (*Lyz2, Il1b*) (B) UMAP showing subclustering analysis of SI-LP lymphoid cluster: NK (*Ncr1, Klrb1c*), NKT cells (*Klrb1c, Cd3d*), innate lymphoid cells type 2 (ILC2, *Il7r, Gata3*) and type 3 (ILC3, *Il7r, Rorc*), γδT-cells (*Cd3d, Trdc*), and 6 clusters of αβT-cells (*Cd3d, Trac*): naïve CD4^+^ (*Sell, Cd4*) and CD8^+^ T-cells (*Sell, CD8a*), mature CD8^+^ T-cells (*Cd44, CD8a*), conventional mature CD4^+^ T-cells (*Cd44, Cd4*), follicular helper CD4^+^ T-cells (*Il21*, T_fh_), Foxp3^+^ regulatory CD4^+^ T-cells (*Foxp3, Il10*, T_reg_) and proliferating cells (*Mki67*). (C) UMAP showing subclustering analysis of SI-LP myeloid cluster: neutrophils (*Itgam, Il1b*), monocytes (*Cx3cr1, Ly6c2*) and macrophages (*Cx3cr1, Apoe*) as wells as 3 populations of dendritic cells: plasmacytoid dendritic cells (pDC, *Siglech, Ly6c2*), conventional dendritic cells type 1 (cDC1, *H2-Ab1, Flt3, Xcr1*) and conventional dendritic cells type 2 (cDC2, *H2-Ab1, Flt3, Itgam, Sirpa, Mgl2*). (D) Plots showing the trends in log_2_ fold-change (FC) of frequencies of respective populations in individual timepoints. Top-left plot shows change in frequencies of 4 major clusters identified in a. Plots on right show changes in frequencies of lymphoid populations from all cells of the lymphoid cluster identified in b. Left-bottom plot shows changes in frequencies of myeloid populations from all cells of the myeloid cluster identified in (C). (E) Scatter plots showing ln FC of average gene expression in lymphoid cluster. Cells from each timepoint (from top to bottom 3-, 4-, 5- and 6-week) are compared to the average expression inside this cluster across all the timepoints. Selected downregulated (blue) and upregulated (red) genes are highlighted. (F) Violin plots of canonical T_H_ subset population markers in conventional CD4^+^ T-cell cluster identified in (A). (G) Violin plots of intraepithelial lymphocytes markers in conventional CD4^+^ T-cell cluster identified in (A).

Having determined the diversity of LP immune cells, we next analyzed changes in the relative abundance of the individual subsets throughout the period of 3-6 weeks of mice age. The most obvious phenomenon was the gradual increase in the frequency of plasma cells (Figure 1D, top-left panel), the majority of which expressed IgA antibody isotype (encoded by the *Igha* gene) (Figure S1A). Next, we observed clear enrichment of myeloid cells and innate lymphocytes peaking at week 4. After this timepoint, the proportion of the innate cell types declined, except for neutrophils that increased in number as late as week 6 (Figure 1D, bottom panels). Among αβT-cells, we observed a gradual decline of naïve subsets associated with their replacement by their mature counterparts (Figure 1D, top-right panel). Collectively, these results suggested that after weaning, mice undergo progressive immune activation in their intestine, with the innate immune branch peaking at 4 weeks of age (i.e. right after weaning) and the adaptive one gradually increasing from 3 to 6 weeks of age.

Based on this data, we next sought to further elucidate what type of adaptive immune response was dominant at particular timepoint. To this end, we analyzed the overall expression profile of all cells of the lymphoid cluster at each timepoint and compared it to the average expression across all timepoints. While lymphoid cells from LP of 3- and 4-week-old mice did not show enrichment of any particular type of adaptive immune response, we observed the upregulation of cytotoxic effector genes like granzymes (*Gzma, Gzmb*) in week 5 and effector T_H_17 cytokines (*Il17a, Il22*) in week 6 (Figure 1E). To gain further insights we specifically focused on cytokines and effector molecules defining major T_H_ subsets. Therefore, we analyzed the expression of interferon gamma (IFNγ, *Ifng*), interleukin-4 (IL-4, *Il4*), IL-17a (*Il17a*) and IL-22 (*Il22*) cytokines typical for T_H_1, T_H_2 and T_H_17 responses respectively. While *Il4* expression was low throughout all the timepoints, *Ifng* showed bimodal expression peaking at weeks 3 and 6. The intestinal expression of *Ifng* at week 3 was reported previously(Al Nabhani *et al*., 2019), however, its gradual increase in later timepoints was not expected. In contrast, T_H_17 effector cytokines (*Il17a* and *Il22*) were expressed virtually only at 6 weeks of age (Figure 1F), potentially explaining the observed accumulation of neutrophils during this time-point (Figure 1D, bottom-left panel). Importantly, conventional CD4^+^ αβT-cells showed an increased expression of molecules necessary for chemotaxis (*Ccr9*) and adhesion (*Itgae, Itgb7*) to IECs from 5 weeks. This phenomenon was also matched by the upregulation of *Cd8a, Gzma* and *Gzmb*, markers typical for IELs (Cheroutre *et al*., 2011; Hayday *et al*., 2001) (Figure 1G).

### Age-dependent upregulation of MHCII and antigen presentation machinery by IECs

To elucidate the potential impact of age-dependent intestinal colonization on the epithelial compartment we performed scRNA-seq of IECs from the same time-points as described above. The clustering analysis highlighted all major IECs populations (Figures 2A and S2) that underwent individual subclustering (Figures S3 and S4). Firstly, we analyzed the changes in relative abundances of IECs throughout the period of 3-6 weeks of mice age. The most pronounced was the continuous increase in Tuft cells and the concomitant decrease in Paneth cells frequencies (Figure 2B). Furthermore, transcriptional analysis of the individual IEC subsets revealed several striking differences that depend on age of the animal including genes encoding MHCII (*H2-Ab1, Cd74*) that were not expressed in IECs during earlier timepoints and became upregulated only at 5 and 6 weeks of age. This suggested that their enhanced expression might be the outcome of immune reaction to changes associated with weaning. In addition to the upregulation of MHCII encoding genes, we also observed a gradual increase in the expression of secretory IgA receptor (*Pigr*) and antimicrobial peptides of the Reg3 family (*Reg3b* and *Reg3g*) (Figures 2C-D) likely induced by IL-17 and IL-22 cytokines from T_H_17 cells (Kumar *et al*., 2016; Shih *et al*., 2014).

**Figure 2.**
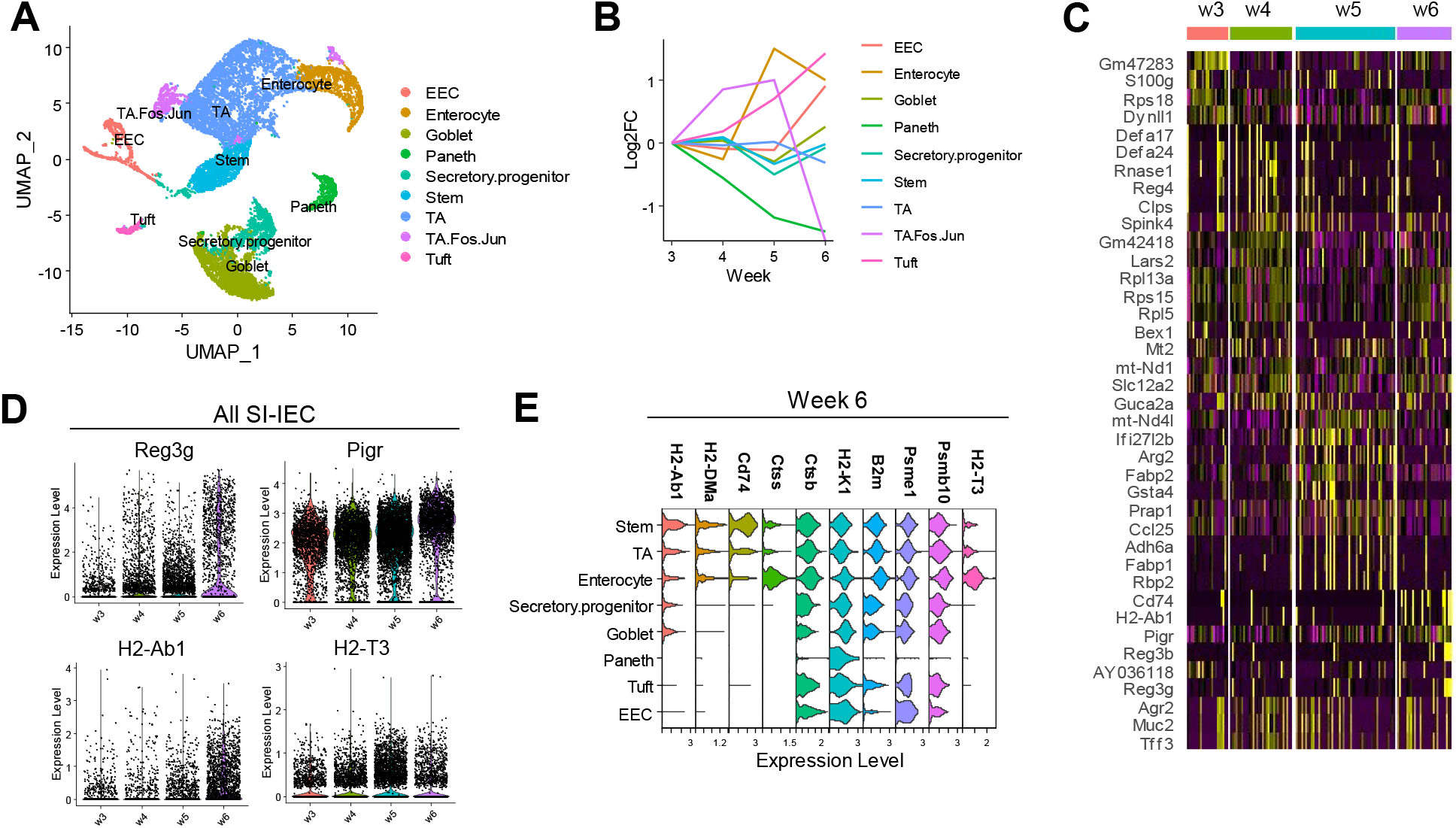
Profiling of small intestinal epithelial cells through mouse age. (A-E) scRNA-seq analysis of SI-IEC from 3-, 4-, 5- and 6-weeks old WT littermate mice. (A) UMAP of IEC clustering analysis. Stem cells (defined based on *Lgr5, Olfm4* expression), transiently amplifying cells (TA, *Mki67, Top2a*), mature enterocytes (*Fabp1, Fabp6*), Enteroendocrine cells (EEC, *Chga, Chgb*), Tuft cells (*Avil, Dclk1*), Paneth cells (*Lyz1, Defa24*) and goblet cells (*Muc2, Tff3*). (B) Plots showing Log2 FC of IEC populations described as frequencies across timepoints. (C) Heatmap showing top 10 differentially expressed genes across all the timepoints in all IECs subsets. (D) Violin plots showing the expression of selected differentially expressed genes across all timepoints in all IECs. (E) Stacked violin plot showing the expression of MHCII (*H2-Ab1, H2-DMa*), cathepsins responsible for MHCII loading (*Ctss, Ctsb*), MHCI (*H2-K1, B2m*), proteasomes responsible for MHCI loading (*Psme1, Psmb10*) and nonclassical MHC molecule TLA (*H2-T3*) in individual IEC populations at 6-weeks of ages.

To determine which of the individual IEC sub-populations acquired the highest antigen presentation potential, we analyzed the expression of MHCII encoding genes (*H2-Ab1, H2-DMa, Cd74*), MHCI encoding genes (*H2-K1, B2m*) together with their respective antigen processing machinery (MHCII: *Ctss, Ctsb*; MHCI: *Psme1, Psmb10)* as well as major CD8αα ligand TLA (*H2-T3)* at 6 weeks, where MHCII expression was the highest. MHCII-associated genes showed the highest expression in the stemcells compartment but were also detectable in Transiently amplifying cells (TA) and enterocytes. Although all these cells also expressed MHCII-associated antigen processing cathepsins, their highest expression was restricted to mature enterocytes. Enterocytes also showed the highest expression of MHCI, as well as proteasome subunits generating substrates for MHCI loading, together with H2-T3 (Figures 2E and S5A-B). Notably, stem cells, TA and enterocytes did not show robust expression of any costimulatory molecules implicated in the priming or activation of the CD4^+^ T-cells (Figure S5C).

A more detailed analysis of intestinal stem cell and TA clusters revealed substantial diversity in the intestinal stem cell compartment, which segregated into two major subsets. This was in line with previous studies, which showed that intestinal stem cells increase their differentiation state with concomitant loss of MHCII expression (Biton *et al*., 2018). Indeed, the first subset highlighted by our analysis consisted of resting stem cells (likely encompassing intestinal stem cells type I and II subsets defined previously (Biton *et al*., 2018)), which expressed high MHCII levels, while the second subset showed high expression of proliferation markers but lower MHCII expression (representing intestinal stem cells type III defined previously (Biton *et al*., 2018)). Interestingly, MHCII expression in all those subsets showed a strict dependence on the age of the mouse, being expressed at high levels only in stem cells from 6-weeks-old mice (Figures S3D-F).

Since enterocytes also upregulated the expression of MHCII encoding genes, we sought to further elucidate the internal diversity of this subset. We first defined populations of enterocytes originating from the duodenum, jejunum and ileum regions (Figure S4A), using previously defined markers (Haber et al., 2017) (Figure S4B). We observed an increasing trend in the MHCII expression from the duodenum to the ileum. This increasing trend was also matched by the expression of cathepsins (Figure S4C). We then subclustered enterocytes from individual small intestinal segments using markers defining the vertical position of the enterocytes on the villus (Moor et al., 2018) (Figures S4D-I). We assessed the expression of antigen presentation machinery and associated genes in defined enterocyte subclusters. Duodenal enterocytes showed almost no MHCII expression (Figures S4C and S4J). Jejunal enterocytes showed a decreasing trend of MHCII expression from the bottom to the top of the villus (Figures S4C and S4K), likely extending the trend of lowering MHCII expression with a higher differentiation state observed among stem cell and TA clusters (Figures S3D-F). In contrast, ileal enterocytes expressed comparable levels of MHCII in all the villus zones (Figure S4L), suggesting that in this specific region MHCII might play an additional role in mature enterocytes biology.

In summary, our data suggest that after mouse weaning, several waves of immune responses are mounted in the SI-LP. First, innate immune cells accumulate at 4 weeks, but are soon replaced by the cells of adaptive immunity from 5 weeks and peaking at 6 weeks of age. We mostly observed T_H_1 and T_H_17 responses, accompanied by an increase in IEL markers expression in CD4^+^ conventional αβTCR T-cells of SI-LP. Adaptive immune responses were matched by the concomitant increase in the expression of MHCII on stem, TA and enterocyte populations of IECs.

### SFB induce strong MHCII expression on IECs

To validate the age-dependent MHCII expression on IECs highlighted by our scRNA-seq analysis, we analyzed the expression of MHCII on IECs (CD45^−^ EpCAM^+^) isolated from 2-7 weeks-old WT littermate mice using flow cytometry. Similar to the scRNA-seq data, we observed that while almost no MHCII was expressed in earlier timepoints (week 2-4 of age), IECs showed remarkable MHCII expression starting from 5 weeks of age (Figures S6A-B). This was further confirmed by immunofluorescent microscopy of the small intestine from 3- and 5-week-old mice, which also confirmed the expression of MHCII on the higher zones of the mature enterocytes’ villi (Figure S6C).

During the routine screening, we noted that genetically identical WT mice housed in different compartments of the local animal facility showed high differences in their expression of MHCII on IECs, defining MHCII-IEC^Low^ and MHCII-IEC^High^ WT strains (Figure 3A). This observation pointed to the possible involvement of environmental factors capable of inducing MHCII^+^ IECs. We transferred a breeding pair of MHCII-IEC^Low^ mice on the bedding from MHCII-IEC^High^ mice. This manipulation alone induced MHCII expression on IECs in the progeny of formerly MHCII-IEC^Low^ parents (Figure 3A), suggesting that MHCII expression on IECs is induced by microbiota. Therefore, to address whether the observed effect is linked to a specific bacterial strain, we next performed 16S-sequencing of ileum microbiota present in MHCII-IEC^Low^ and MHCII-IEC^High^ mice. While the MHCII-IEC^Low^ animals had a higher abundance of *Dorea* and *Ruminicoccus* taxa (Figure 3B), the MHCII-IEC^High^ mice showed a higher abundance of *Lactobacillus, Blautia* taxa and *Candidatus arthromitus*, also known as SFB, which was previously suggested to modulate MHCII expression on the intestinal epithelium (Goto et al., 2014; Tuganbaev *et al*., 2020; Umesaki et al., 1995).

**Figure 3.**
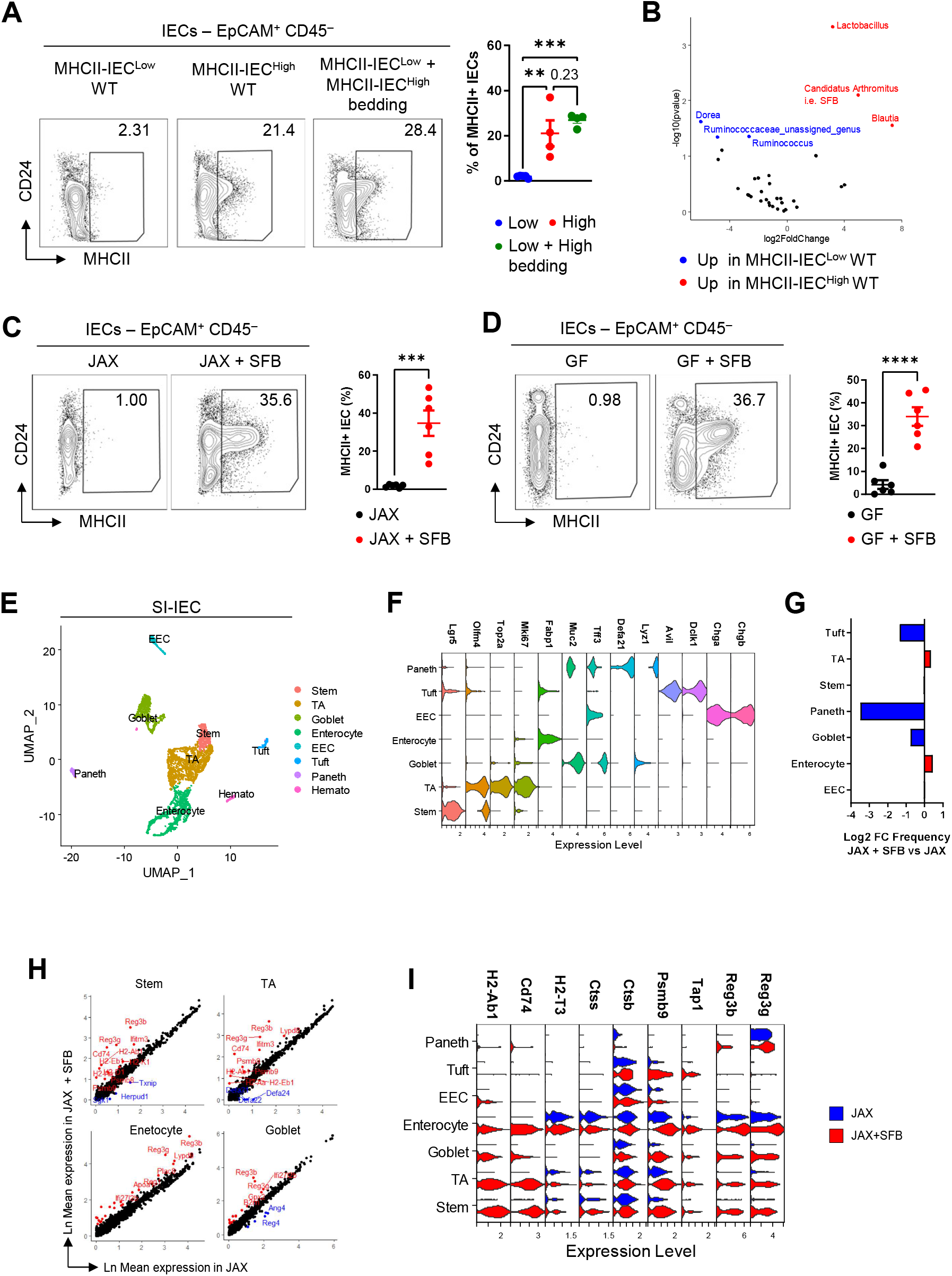
SFB induce MHCII and antigen processing machinery in intestinal epithelial cells. (A) FACS analysis of SI-IECs isolated from 5-week-old WT mice from two separately housed colonies (MHCII-IEC^Low^ WT and MHCII-IEC^High^ WT) and the progeny of MHCII-IEC^Low^ WT breeding pair, which was housed on the bedding from MHCII-IEC^High^ WT mice (MHCII-IEC^Low^ WT + MHCII-IEC^High^ WT bedding). FACS plots show representative gating. Numbers beside gates show frequency from parent population. Plot on the right shows overview of the results and statistical analysis (n=4-5). (B) Volcano plot of 16S sequencing analysis of ileum microbiota from MHCII-IEC^Low^ WT and MHCII-IEC^High^ WT animals. Selected taxa with higher abundance in MHCII-IEC^Low^ WT (blue) and MHCII-IEC^High^ (red) are highlighted. Data was tested by DESeq2 and shown as -Log10 of plain pvalue compared to Log2 FC. n=3. (C) FACS analysis of SI-IECs isolated from 5-week-old WT animals carrying microbiota from Jackson Laboratories (JAX), half of which was colonized with SFB at 21 and 22 days of age. Left FACS plots show representative gating. Plot on the right shows overview of the results and statistical analysis. Numbers beside gates show frequency from parent population. n=6. (D) FACS analysis of SI-IECs isolated from 5-week-old germ-free (GF) WT animals, half of which was colonized with SFB at 21 and 22 days of age. Left plots show representative gating. Plot on the right shows overview of the results and statistical analysis. Numbers beside gates show frequency from parent population. n=6. (E-I) scRNA-seq analysis of SI-IEC from 5-weeks-old WT JAX mouse and a littermate, which was colonized with SFB at 21 and 22 days of age. (E) UMAP of IEC clustering analysis. (F) Stacked violin plot showing the expression of canonic marker genes, defining the annotation of cell types. (G) Bar graph showing Log2 FC of frequency of IEC populations comparing SFB colonized to noncolonized mice. (H) Scatter plots showing Ln FC of average gene expression in respective populations, comparing SFB colonized to noncolonized mice. Selected downregulated (blue) and upregulated (red) genes are highlighted. (I) Stacked violin plot comparing the expression of selected differentially expressed genes between SFB colonized and noncolonized mice across all IEC populations. Horizontal lines in (A) (C) and (D) show mean±SEM. Data in (A) was tested by Ordinary one-way ANOVA with Holm-Šídák’s multiple comparisons test. Data in (C) and (D) was tested by Student’s t-test. **, p<0.01; ***, p<0.001; ****, p<0.0001; p values >0.05 are shown.

To confirm whether SFB can indeed induce MHCII expression on IECs, we colonized WT mice acquired from Jackson laboratories vendor (JAX) by SFB, as mice acquired from JAX were previously shown to be SFB-free (Ivanov *et al*., 2009) (Figure 3C). Furthermore, we performed SFB mono-association of germ-free animals (Figure 3D). SFB induced strong expression of MHCII on IECs in both settings, suggesting that MHCII expression on IECs is a hallmark of the SFB induced response and that it is independent of the presence of other bacteria.

Having identified SFB as the potent inducers of MHCII expression on IECs, we sought to define its effect on the IEC compartment in more detail. Thus, we performed scRNA-seq of IECs derived from JAX mice or their littermates, colonized with SFB on top of their standard microbiota. Similar to previous experiments, we first identified all the canonic IEC populations in our dataset (Figures 3E-F). We then compared the composition of the populations in SFB colonized and non-colonized mice. We observed a large drop of Paneth cells and to a lower extent also Tuft cell frequency (Figure 3G). We confirmed the decrease of Paneth cell frequency using flow cytometry (Figure S7A). Next, we compared the transcriptome of JAX and SFB-colonized IEC populations. Strikingly, among the most differentially expressed genes were MHCII, together with its antigen processing machinery and TLA (*H2-T3*) molecule. This was accompanied by the induction of Reg3 (i.e. *Reg3b* and *Reg3g*) proteins in the presence of SFB (Figures 3H-I). Notably, Reg3 proteins were previously shown to prevent SFB overgrowth in an IL-22 dependent manner (Shih *et al*., 2014).

Analogically to previous experiments, we also subclustered EECs (Figures S7B-D), stem cells (Figures S7E-H) and mature enterocytes (Figure S8). Among stem cell subtypes, we observed an increase in the frequency of the resting stem cells, matched by the drop in the frequency of proliferating ones (Figures S7E-G). MHCII expression of all the stem cell subtypes showed strict dependence on the SFB colonization (Figure S7H). Lastly, we subclustered mature enterocytes according to their origin from the duodenum, jejunum, or ileum as well as according to their vertical position on the villus (Figure S8). MHCII expression in SFB colonized mice showed an increasing trend from the duodenum to the ileum. Moreover, it was expressed in high levels not only by bottom zone enterocytes but also by the ones residing in higher zones of the villus in both the duodenum and ileum (Figure S8).

Collectively, these results show that mice colonized with SFB phenotypically match 6-week old mice (Figure 1) and identify SFB as a key inducer of MHCII and antigen presentation machinery expression in different IEC subsets, including mature enterocytes.

### MHCII expression on IEC is induced by T-cell-derived IFNγ in a cDC1-dependent manner

While SFB are best known for their induction of a T_H_17 response (Gaboriau-Routhiau *et al*., 2009; Ivanov *et al*., 2009), it was also reported that SFB can increase IFNγ production in the gut (Gaboriau-Routhiau *et al*., 2009). The latter finding may be crucial since IFNγ is a well-documented modulator of MHCII expression (Koyama *et al*., 2019; Thelemann *et al*., 2014; Van Der Kraak *et al*., 2021). Therefore, we next measured the induction of IL-17a and IFNγ cytokines in the SI-LP after SFB colonization of JAX mice. As expected, SFB induced a strong Th17 responses in the SI-LP. Although the frequency of IFNγ^+^ T_H_1 cells was not significantly changed, we observed a change in the magnitude of IFNγ expression by T_H_1 (Figure 4A). We did not detect any major differences in the IFNγ or IL-17 production in CD8^+^ SI-LP T-cells, or IELs (Figure S9).

**Figure 4.**
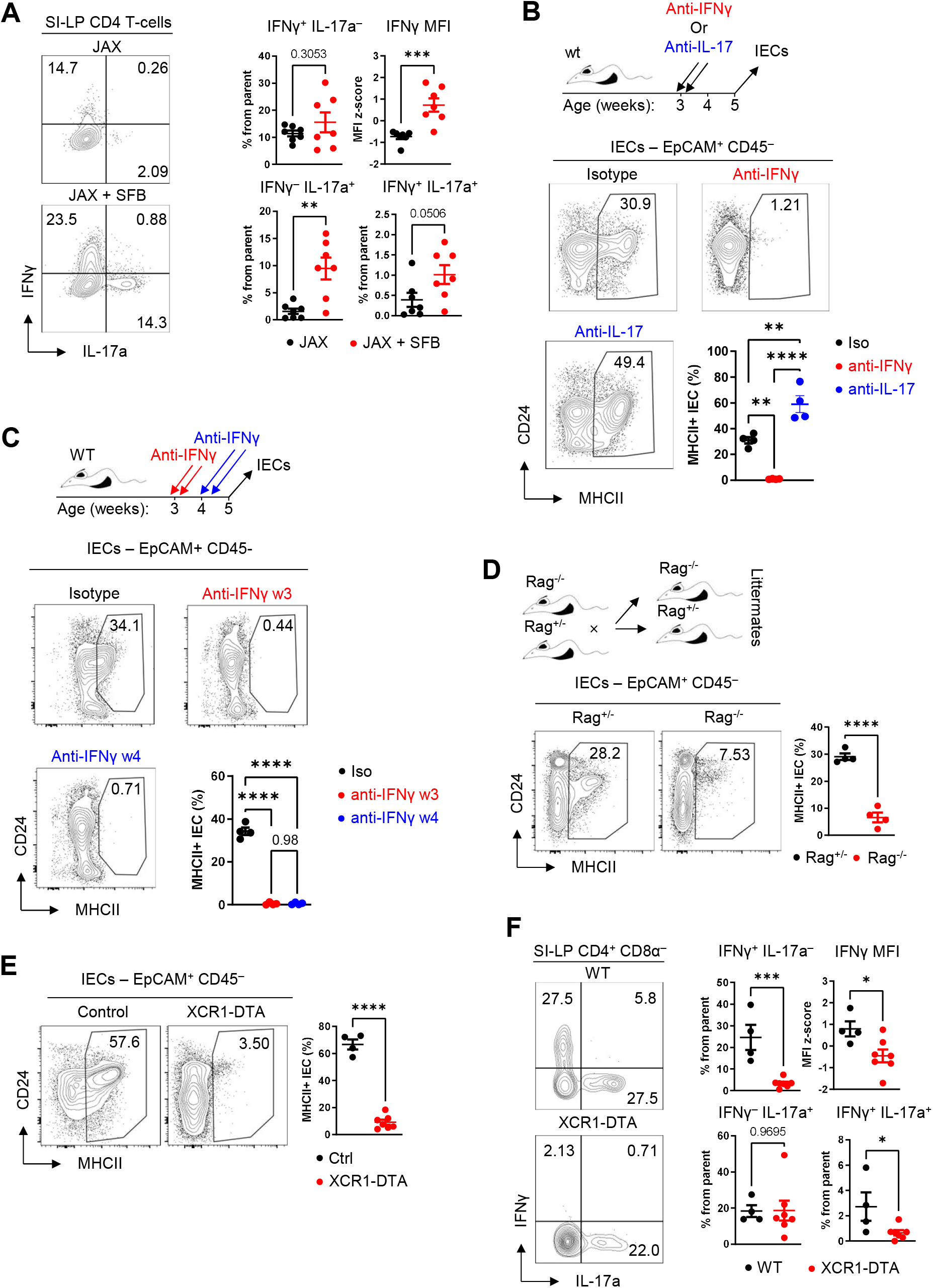
cDC1-controlled T-cells regulate MHCII expression on IECs through the production of IFNγ. (A) Flow cytometry analysis of IL-17 and IFNγ expression in SI-LP CD4^+^ T-cells in 5-weeks-old JAX mice, and their SFB-colonized littermates. FACS plots on the left show representative gating. Plots on the right show overview of frequencies of cytokine positive T-cells and the z-score of IFNγ median fluorescence intensity (MFI). n=7. (B) Flow cytometry analysis of MHCII expression on SI-IECs in SFB colonized mice, which were i.p. treated with either IL-17 or IFNγ neutralizing antibodies or isotype control at 21 and 22 days of age, 250 µg in 100 µl of PBS in each injection. Plots show representative gating and the statistical overview of the data. n=4. (C) WT SFB colonized mice were i.p. treated either with isotype antibody or with IFNγ neutralizing antibody at either 21 and 22 days or at 28 and 29 days of age, 250 µg in 100 µl of PBS in each injection. Plots show representative gating and statistical overview of the data. Numbers beside gates represent frequencies. n=4. (D) Flow cytometry analysis of MHCII expression on SI-IECs in SFB colonized Rag1+/- and Rag1-/-littermate mice at 5 weeks of age. n=4. (E) Flow cytometry analysis of MHCII expression on SI-IECs in SFB colonized XCR1-Cre x R26-fl-STOP-fl-DTA (XCR1-DTA) and R26-fl-STOP-fl-DTA (control) littermate mice at 5 weeks of age. n=4-7. (F) Flow cytometry analysis of IL-17 and IFNγ expression in SI-LP C4+ T-cells in 5-weeks-old XCR1-DTA and control littermate mice. FACS plots on the left show representative gating. Plots on the right show overview of frequencies of cytokine positive T-cells and the z-score of IFNγ MFI. n=4-7. Horizontal lines show mean±SEM. Data in (A) and (D-F) was tested by was tested by Student’s t-test. Data in (B) and (C) was tested by Ordinary one-way ANOVA with Holm-Šídák’s multiple comparisons test. *, p<0.05; **, p<0.01; ***, p<0.001; ****, p<0.0001; p values >0.05 are shown.

To test, whether SFB-induced IFNγ or IL-17 affect MHCII expression on IECs, we blocked the effector function of these two cytokines in SFB-colonized littermate mice by neutralizing antibodies. Strikingly, while IFNγ blockade completely abrogated MHCII expression on IECs, blockade of IL-17 led to a2-foldld increase in the frequency of MHCII^+^ IECs (Figure 4B). Since a recent study reported that during weaning, mice show strong but transient production of IFNγ (Al Nabhani *et al*., 2019), we wondered whether blockade of this transient IFNγ stimulus associated with weaning could affect the MHCII expression on IECs later in life. To this end, we injected WT SFB^+^ mice with IFNγ blocking antibody either at 3 or at 4 weeks of age and analyzed them at the age of 5 weeks. In both cases, the MHCII expression on IECs was completely blocked. Thus, continuous IFNγ production seems to be indispensable for the expression of MHCII on IECs not only during but also after the weaning period (Figure 4C).

Next, we soughed to identify the cellular source of IFNγ responsible for the induction of MHCII expression on IECs. We utilized scRNA-seq analysis of SI-LP from 6-weeks old mice (Figure 1), which showed a robust expression of MHCII on IECs. In this dataset, 3 main populations of SI-LP CD45^+^ cells expressed IFNγ: CD4^+^ αβT-cells, NK cells, and NKT cells (Figure S10A). Using flow cytometry, we found that although NK cell (Nk1.1^+^ CD3ε^−^), NKT cells (Nk1.1^+^ CD3ε^+^), as well as further undefined CD3ε^−^ population indeed produce IFNγ, CD3ε^+^ T-cells are by far the most numerous producers of IFNγ in the SI-LP (Figure S10B).

To determine whether T cells may be the key inducer of the MHCII expression on IECs, we analyzed Rag1-deficient mice that lack mature T-cells. To eliminate the possible effect of litter and/or separate housing we compared Rag1^−/–^ to their Rag1^+/–^ littermates. Using this system, we observed the reduction of MHCII expression on IECs by ∼75% in the absence of mature T-cells (Figure 4D). Next, to test whether NK/NKT may also contribute to the induction of MHCII expression on IECs, we depleted NK and NKT cells using NK1.1 antibody in SFB^+^ WT mice, based on a previously described protocol (Glasner et al., 2018). Although this treatment largely depleted both NK cells (by ∼85%) and NKT cells (by ∼50%) (Figure S10C), we observed no significant effect on the frequency of MHCII^+^ IECs (Figure S10D). Suggesting, that IFNγ-producing T-cells are the major driver of MHCII expression on IECs in lympho-sufficient mice.

Since T-cell responses are canonically driven by specific antigen presenting cells (APCs) populations, we next wanted to determine if SFB-induced IFNγ producing T-cells, responsible for the induction of MHCII on IECs are also regulated in such a way. Xcr1^+^ cDC1 were recently identified as the major inducers of T_H_1 responses after intestinal infection by epithelial-adhesive cryptosporidium (Russler-Germain et al., 2021). To test whether cDC1 may also be involved in the SFB-mediated induction of MHCII^+^ on IECs, we next utilized and analyzed SFB^+^ Xcr1-Cre x R26-flox-STOP-flox-DTA (Xcr1-DTA) mice (Voehringer et al., 2008; Wohn et al., 2020), in which the mature cDC1 population is deleted by inducible diphtheria toxin expression (Figure S11A). Strikingly, these mice showed a ∼95% decrease in the MHCII expression on their IECs compared to their cDC1-sufficient littermates (Figure 4E). Furthermore, they almost completely lacked IFNγ expression in both CD4^+^ and CD8^+^ SI-LP T-cells, while IL-17 production by CD4^+^ LP T-cells was unaffected (Figures. 4F and S11B). Collectively, these results suggest that MHCII on IECs is induced by IFNγ that is produced by T-cells under the control of the cDC1 professional APCs population.

### SFB induce granzymes-expressing iIELs in the small intestine

While SFB-induced Th17 responses were studied extensively, other immune responses elicited by this microbe remain rather poorly described. To get a broader perspective of immune responses directed against SFB, we performed scRNA-seq of SI-LP cells isolated either from SFB-free or SFB-colonized JAX mice. The clustering analysis highlighted several major subsets, including B-cells, plasma cells, several types of myeloid cells, ILC2 and 3 cells and a cluster containing other lymphoid cells. Further analysis of the mixed lymphoid cluster highlighted several sub-clusters corresponding to NK cells, NKT cells, both naïve and mature γδT-cells, naïve and mature CD8^+^ αβT-cells, CD4^+^ naïve T-cells, T_reg_s and conventional CD4^+^ T-cells (Figures 5A and S12A).

**Figure 5.**
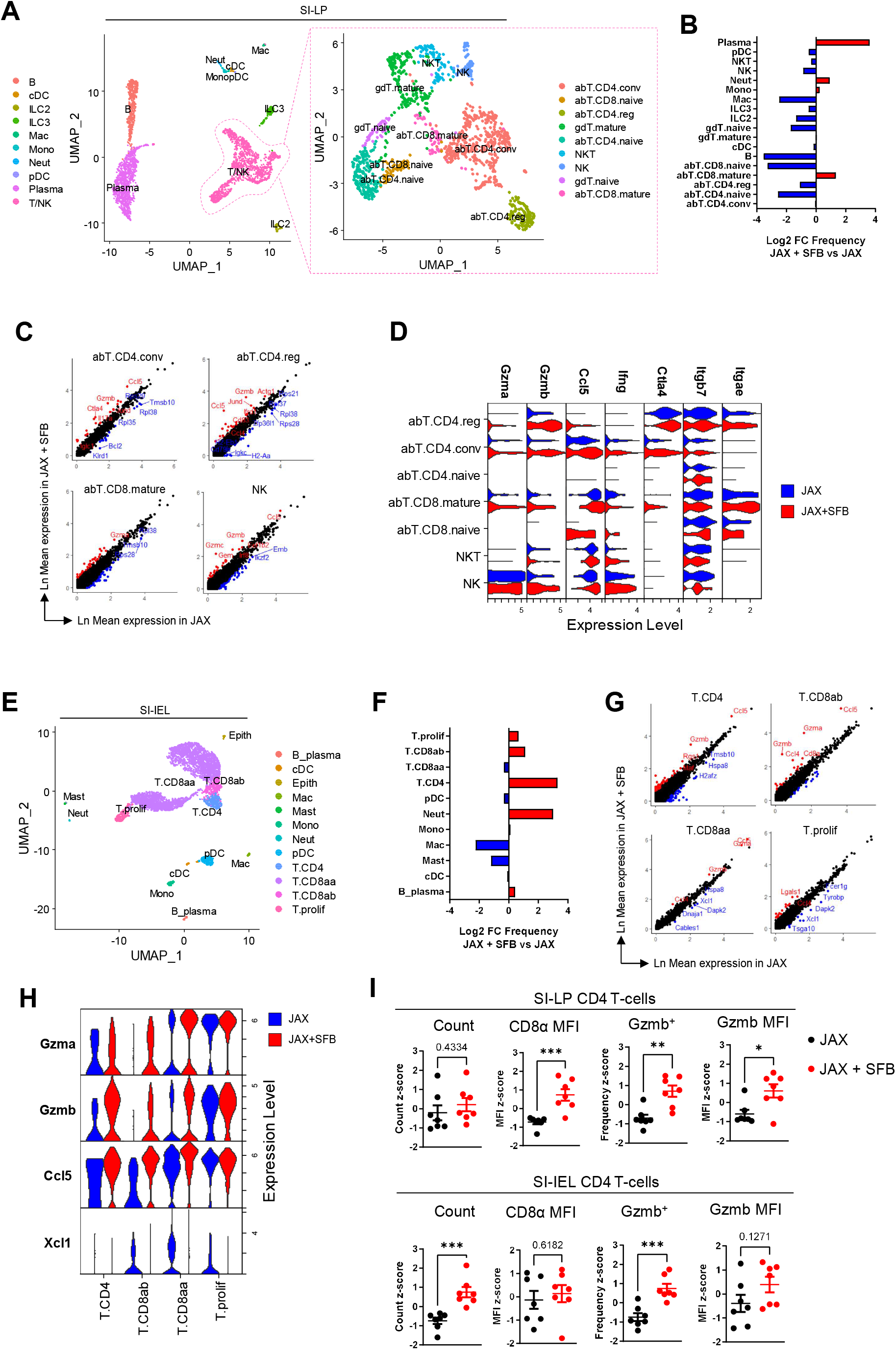
SFB induce Gzmb+ intraepithelial CD4+ T cell responses. (A-D) scRNA-seq analysis of SI-LP cells from JAX and SFB colonized JAX mice. (A) UMAP on the left shows basic clustering to major cell types. UMAP on the right shows subclustering of T/NK cluster. (B) Bar graph showing Log2 FC of frequency of SI-LP populations comparing SFB colonized to noncolonized mice. (C) Scatter plots showing Ln FC of average gene expression in respective populations, comparing SFB colonized to noncolonized mice. Selected downregulated (blue) and upregulated (red) genes are highlighted. (D) Stacked violin plot comparing the expression of the selected differentially expressed genes between SFB colonized and noncolonized mice across all SI-LP T- and NK cell populations. (E-H) scRNA-seq analysis of SI-IEL cells from JAX and SFB colonized JAX mice. (E) UMAP showing clustering and cell type annotation of SI-IEL cells. (F) Bar graph showing Log2 FC of frequency of SI-IEL populations comparing SFB colonized JAX mice to to noncolonized littermates. (G) Scatter plots showing Ln FC of average gene expression in respective populations, comparing SFB colonized to noncolonized mice. Selected downregulated (blue) and upregulated (red) genes are highlighted. (H) Stacked violin plot comparing the expression of the most changed genes between SFB colonized and noncolonized mice across all SI-IEL T-cell populations. (I) Flow cytometric analysis of SI-LP (top row) and SI-IEL (bottom row) CD4^+^ T-cells from JAX and SFB colonized JAX mice. Data are batch-normalized using z-score. Plots show (from left): Overall cell count, MFI of CD8α, frequency of Gzmb^+^ cells, MFI of Gzmb. n=7. Horizontal lines in (I) show mean±SEM. Data in (I) was tested by Student’s t-test. *, p<0.05; **, p<0.01; ***, p<0.001; p values >0.05 are shown.

Next, we analyzed changes in the frequencies of SI-LP cell populations after SFB colonization. Most striking was the massive increase in the frequency of plasma cells, which was matched by the drop in the frequency of naïve B-cells. Similarly, we observed a drop in naïve T-cell populations, which were replaced by their mature counterparts. Among innate immune cells, only neutrophils and monocytes increased in frequency (Figure 5B). These findings are in line with previous publications reporting that SFB promote B-cell maturation in plasma cells, marked by IgA production (Lecuyer *et al*., 2014) as well as IL-17-dependent neutrophil accumulation (Flannigan et al., 2017).

To determine the overall effect of SFB colonization on T-cell-dependent adaptive immune responses, we compared the transcriptome of mature T cell populations derived from JAX WT mice, and their SFB colonized littermates. Strikingly, genes most upregulated by SFB colonization were granzymes (*Gzma, Gzmb*) and *Ccl5*. These genes were induced across all the mature T-cell populations and even in NK cells. As expected, conventional CD4^+^ αβT-cells also increased the expression of *Il17a*. However, the same population also showed the induction of inhibitory *Ctla4* molecules (Figures 5C-D). While granzymes are classically associated with cytotoxic lymphocytes (i.e. CD8^+^ T-cells, NK cells, and NKT cells), their expression is also the hallmark of IELs. Therefore, we also analyzed the expression of integrins necessary for lymphocytes to attach to IECs in order to become IELs (i.e. *Itgb7* and *Itgae*) (Cheroutre *et al*., 2011). We have found that SFB colonization induced the expression of these genes in T_reg_ and conventional CD4^+^ αβT-cells in SI-LP (Figure 5D).

Since our data point to the possibility that SFB induce iIEL response we next performed scRNA-seq of SI-IELs from JAX mice and their SFB-colonized littermates. In this dataset, we defined several populations of T-cells (*Cd3d*): mixed population of proliferating T-cells (*Mki67*), CD8αα^+^ T-cells (*CD8a*, classically termed natural IEL – nIEL), CD8αβ^+^ αβT-cells (*Cd8a, Cd8b*) and CD4^+^ αβT-cells (*Cd4*) (latter two are collectively termed iIEL), which together represented the large majority of analyzed cells. Furthermore, we were able to define several myeloid populations: Mast cells (*Cpa3*), neutrophils (*Il1b*), monocytes (*Lyz2, Ly6c2*), cDCs (*Flt3*), pDCs (*Siglech*) and macrophages (*Apoe*). We also defined small populations of B/plasma cells (*Cd19, Sdc1*) and epithelial cells (Figures 5E and S12B). Strikingly, we found that SFB presence increased the frequency of CD4^+^, and to a lesser extent also CD8αβ^+^ iIEL T-cells (Figure 5F). Furthermore, these cells also showed a strong increase in the expression of granzymes (*Gzma, Gzmb*) after SFB colonization (Figures 5G-H).

These observations were confirmed by flow cytometry analysis of the T-cell compartment in SI-LP and SI-IEL of JAX and SFB colonized mice. We were able to define CD4^+^, CD8αβ^+^ as well as CD8αα^+^ IELs (Figure S13A). Importantly, while the counts of all T-cell populations remained unchanged in SI-LP, all of them increased dramatically in SI-IEL after SFB colonization. In both SI-LP and SI-IEL CD4^+^ T-cells showed increased expression of Gzmb. Furthermore, SI-LP CD4^+^ T-cells showed an increase in the expression of CD8α, suggesting the higher rate of their conversion and migration to intraepithelial space (Figures 5I and S13B-E).

We have shown that aside from canonically studied Th17 response, SFB increase CD4^+^ T-cells transition from SI-LP to intraepithelial space, resulting in the accumulation of IEL. Furthermore, SFB modulate proportions of IEL subsets in favor of CD4^+^ iIEL. On top of that, SFB induce the expression of granzymes in both LP and IEL CD4^+^ T-cells.

### MHCII on IECs is needed for SFB-specific iIELs presence

Next, we asked whether the observed SFB-induced iIELs response might be antigen-specific for SFB. We utilized T-cell receptor (TCR) transgenic mouse strain 7B8, where all the T-cells express TCR specifically recognizing SFB-derived antigen bound to MHCII (SFBtg)(Yang *et al*., 2014). First, we performed a fate mapping experiment in which CD44-negative naïve CD90.1/2 CD4^+^ SFBtg T-cells isolated by magnetic separation were intravenously (i.v.) injected into WT (CD90.2/2) SFB^+^ recipients (Figure 6A). To track these cells, we used the congenic marker CD90.1 allowing a clear distinction between transferred and host T-cells (Figure S14A). Using this system, we analyzed the expression of cytokines and IEL-associated molecules in transferred cells in mesenteric lymph nodes (mLN), Payer’s patches (PP), SI-LP and SI-IEL by flow cytometry. Notably, SFBtg T-cells showed a gradual increase in their IL-17a production from mLN to SI-LP, but this cytokine was virtually absent in IELs (Figure 6B). In contrast, Gzmb and CD8α expressions were mostly enriched in SFBtg T-cells derived from the SI-IEL compartment (Figures 6C-D).

**Figure 6.**
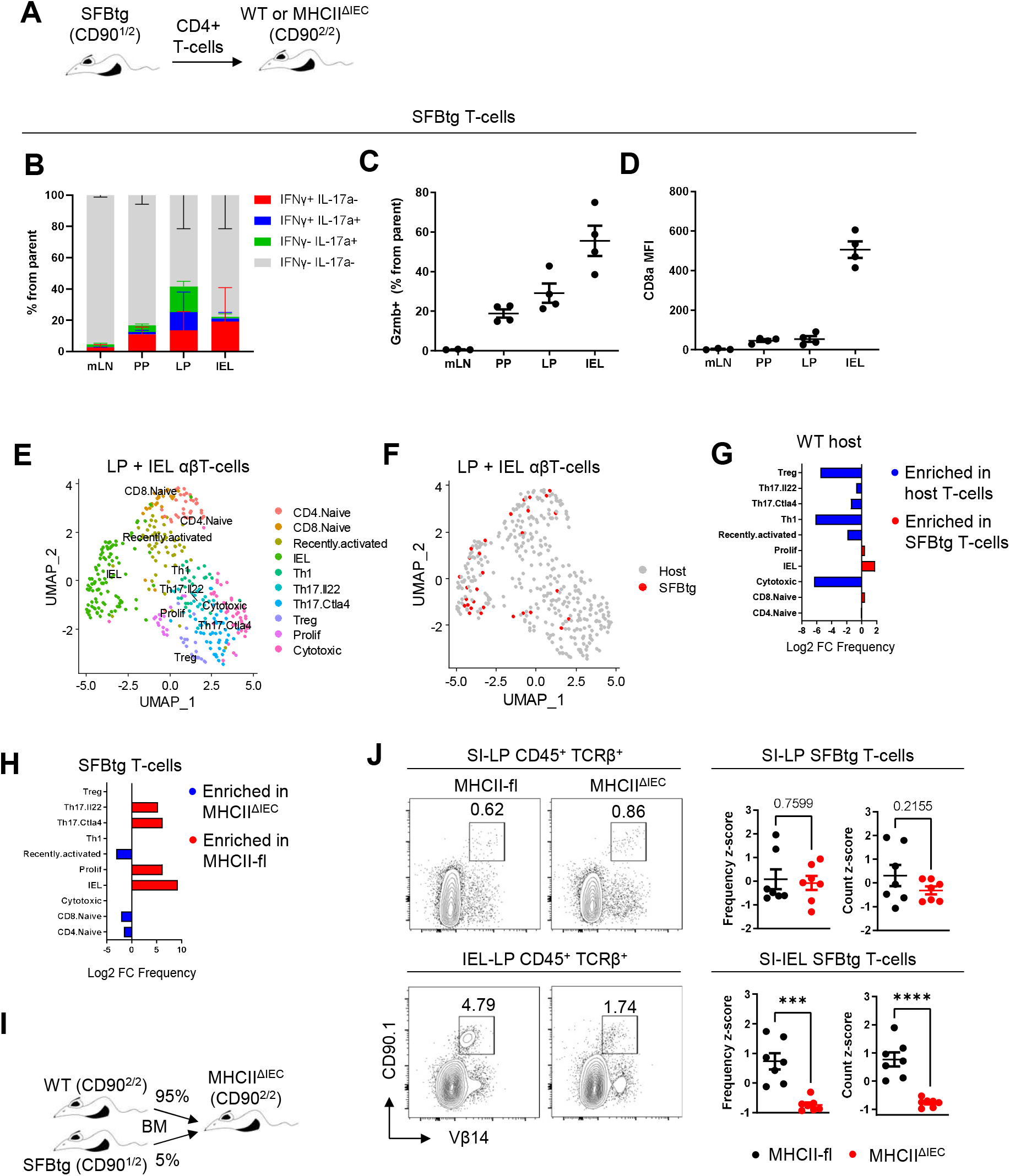
MHCII on IECs controls SFB-specific intraepithelial T-cell responses in the intestine. (A) Schematic showing experimental setup for panels(B-G). TCR-transgenic mice, carrying MHCII-restricted TCR specific for SFB-derived antigen (SFBtg, CD90.1/2) were used as donors of naïve CD4^+^ T-cells. These cells were adoptively transferred to MHCII^ΔIEC^ mice or their cre-negative littermates at 5 weeks of age of recipients. Mice were analyzed 2 weeks after T-cell transfer. (B-D) Flow cytometry analysis of SFBtg T-cell fate after adoptive transfer to WT SFB^+^ host. Transferred cells were analyzed in mesenteric lymph nodes (mLN), Payer’s patechs (PP), SI-LP and SI-IEL. n=4. (B) Relative distribution of IFNγ and IL-17a expressing populations among SFBtg T-cells in indicated tissues. (C) Frequency of Gzmb^+^ cells among SFBtg T-cells in indicated tissues. (D) MFI of CD8α in SFBtg T-cells in indicated tissues. (E-G) Well-based scRNA-seq analysis of SFBtg T-cell fate after adoptive transfer to SFB+ host. Cells were FACS-sorted as TCRβ^+^ live cells from SI-LP and SI-IEL of SFB^+^ MHCII^ΔIEC^ mice and cre-negative littermates (WT). (E) UMAP showing clustering and cell type annotation of pooled SI-IEL and SI-LP cells. (F) UAMP showing origin of T-cells (based on the protein CD90.1 expression) and their affiliation to respective clusters. (G) Bar graphs showing Log2 FC of frequency of individual cell types comparing SFBtg cells to host T-cells. (H) Bar graphs showing Log2 FC of frequency of SFBtg cells distribution among T-cells clusters comparing WT to MHCII^ΔIEC^ hosts. (I) Schematic showing experimental setup for panel (J) Congenically marked SFBtg BM was mixed with feeder WT BM in 5:95 ratio and i.v. injected to irradiated WT or MHCII^ΔIEC^ hosts. MHCII^ΔIEC^ hosts were induced with tamoxifen at 21 and 22 days of age and the BM transplantation was carried out at 4 weeks of age of the acceptors. These mice were analyzed 4-6 weeks after BM transplantation. (J) Flow cytometry analysis of SFBtg T-cell numbers in SI-LP (top row) and SI-IEL (bottom row) of MHCII^ΔIEC^ and littermate WT recipients. FACS plots show representative gating and numbers beside gates show frequency. Plots on the right show z-score batch-normalized statistical overview of the frequency and counts of SFBtg T-cells from all TCRβ^+^ cells in SI-LP and SI-IEL. n=7. Horizontal lines in (B-D) and (J) show mean±SEM. Data in (J) was tested by Student’s t-test. ***, p<0.001; ****, p<0.0001; p values >0.05 are shown.

To assess if SFBtg T-cells differentiate to *bona fide* IELs and to determine the impact of IEC-specific MHCII expression on this process, we performed well-based scRNA-seq of SI-LP and SI-IEL αβT-cells from mice lacking MHCII on their IECs (Villin1-Cre x MHCII-fl/fl: MHCII^ΔIEC^) and their cre-negative littermates harboring adoptively transferred CD90.1^+^ SFBtg naïve T-cells. Index flow cytometry-based sorting allowed us to compare protein expression with subsequent transcriptomic analysis and permitted the distinction of SFBtg T-cells (TCRβ^+^ Vβ14^+^ CD90.1^+^) from host T-cells (TCRβ^+^ CD90.1^−^). In this dataset we were able to define several populations coming from the host SI-LP and SI-IEL T-cells: CD4^+^ (CD4, *Sell, S1pr1, Ccr7*) and CD8^+^ (CD8α, CD8β, *Sell, S1pr1, Ccr7*) naïve T-cells, a compact population of IEL T-cells (*Gzmb, Itgae*), several populations of mature T-cells (*Cd44*) and a population clustering in-between naïve and mature cells, that we designated as recently activated T-cells. Among mature host T-cell populations we found a clearly defined population of T_reg_ cells (*Foxp3, Il10*) and proliferating T-cells (*Mki67, Top2a*), T_H_1 cells (*Ifng*), two populations of T_H_17 cells (*Il17a, Il17f*) defined by opposing trends in the expression of *Ctla4* and *Il22* as well as the further undefined population of cell that expressed high levels of *Gzmb*, which we designated as cytotoxic T-cells (Figures 6E and S14B-D).

Next, we mapped adoptively transferred SFBtg T-cells on the clustering map of the host-derived T-cells. Strikingly, the majority of SFBtg T-cells fell into the IEL cluster (Figure 6F). This data proves that SFBtg IEL cells are transcriptionally indistinguishable from canonical IELs and thus can be designated as *bona fide* members of this population. We then focused on the preferential differentiation states, taken by SFBtg T-cells. The comparison with host-derived T-cells showed that SFBtg T-cells preferentially differentiate into an IEL population, but are underrepresented in T_reg_, T_H_1 and cytotoxic populations (Figure 6G). Notably, the differentiation of SFBtg T-cells into IEL, but also to T_H_17, is highly dependent on the expression of MHCII by IECs (Figure 6H).

Although these experiments allowed for well-controlled fate mapping, their major limitation is the analysis of a particular timepoint after SFBtg T-cells injection, thus we could not exclude the possibility, that the effect of MHCII deletion on IECs could lead to the different kinetics in SFBtg T-cells differentiation, migration or retention in the intraepithelial space in MHCII^ΔIEC^ mice. To overcome this, we constructed bone marrow (BM) chimeric animals, where CD90.2/2 MHCII^ΔIEC^ mice and their cre-negative littermates served as recipients and 5% of CD90.1/2 SFBtg mice mixed with 95% of WT CD90.2/2 feeder cells served as BM donors (Figure 6I). In BM chimeras, SFBtg T-cells are continuously replenished from engrafted BM and thus the observed effect should not be the result of different time kinetics of T-cell differentiation. Strikingly, in MHCII^ΔIEC^ mice SFBtg T-cells were decreased by ∼80% in both frequencies and counts in the intraepithelial space, compared to their cre-littermates (Figure 6J), suggesting that MHCII expression on IECs is crucial for the proper antigen-specific IEL response to SFB. We also analyzed both the WT and SFBtg T-cells cytokine and granzyme expression in SI-LP and SI-IEL. While SFBtg T-cells in the SI-LP showed a drop in IFNγ^+^ IL-17^+^ population as well as their Gzmb expression, their IELs counterparts did not show any clear differences in the expression of these molecules (Figures S15A and S15C). However, it should be noted that very few SFBtg T-cells are left in the IEL compartment of MHCII^ΔIEC^ mice (Figure 6J). Among WT T-cells, the only population that was affected were SI-LP IFNγ^+^ CD4^+^ T-cells, whose frequency was slightly decreased (Figure S15B).

In summary, this data demonstrates that SFB presence predispose SFB-specific CD4^+^ T-cell clones to localize into the intraepithelial space and differentiate into Gzmb^+^ iIELs. Moreover, we show that this process is largely dependent on MHCII expression on IECs.

### MHCII on the intestinal epithelium and hematopoietic granzymes regulate the epithelial turnover

Finally, we sought to elucidate the putative role of SFB-induced Gzmb^+^ iIELs and their potential impact on the intestinal epithelium physiology.

Therefore, we next aimed to test whether the epithelial expression of MHCII in SFB colonized animals could regulate the epithelial turnover. To this end, we injected MHCII^ΔIEC^ and their cre-negative littermates with 5-ethynyl-2’-deoxyuridine (EdU) that incorporates to newly synthesized DNA and analyzed the amount of EdU^+^ IECs 72 hours later (Figure 7A), a time necessary for IEC to travel from crypt to the villus tip. Strikingly, we detected a ∼1.5-fold increase of EdU^+^ IECs in MHCII^ΔIEC^ mice using both flow cytometry (Figure 7B) and microscopic analysis of intestinal sections (Figure 7C), suggesting MHCII on IECs promotes IECs trafficking to villus tip and some of them are shed into SI lumen in a shorter timescale than 72 hours.

**Figure 7.**
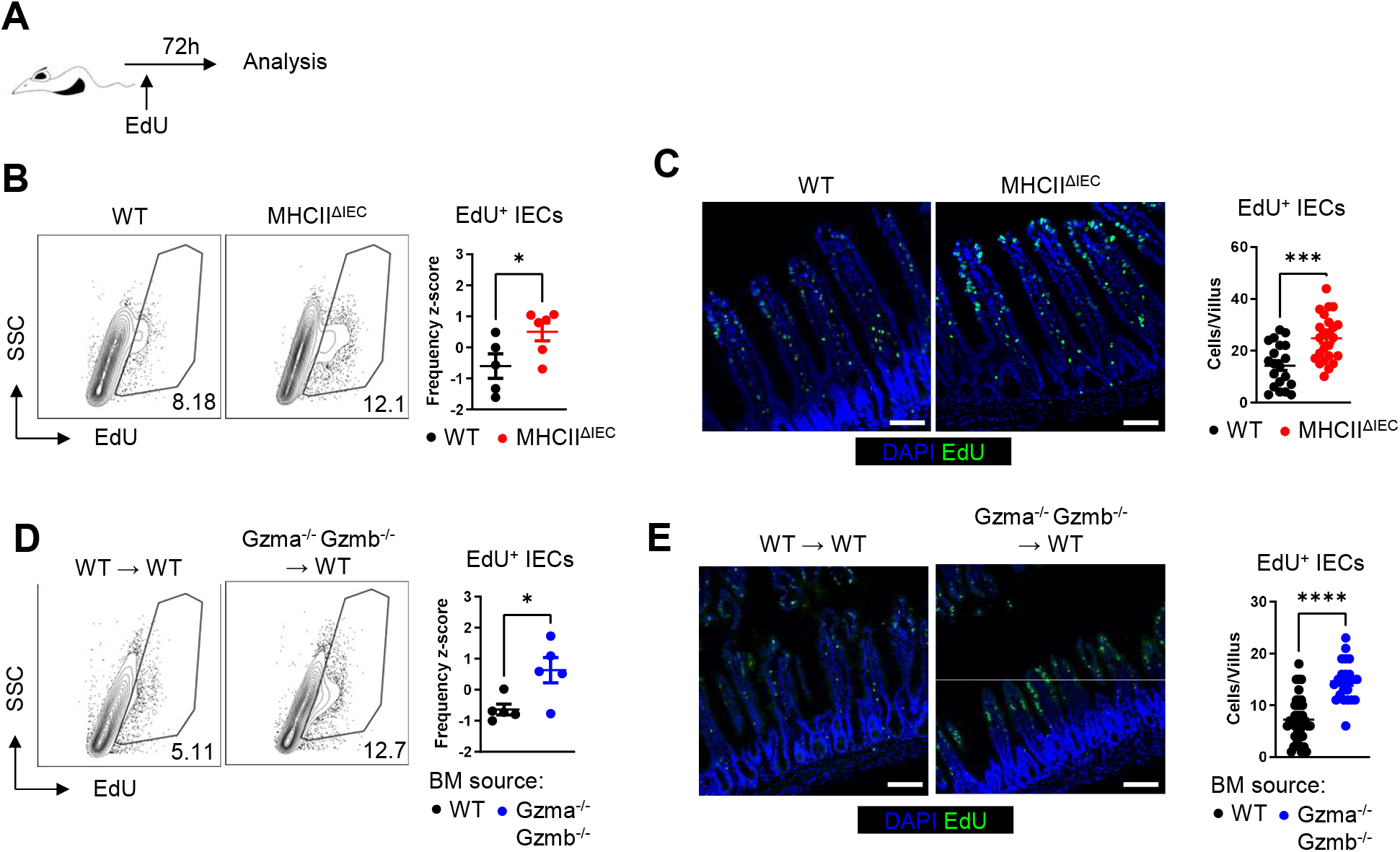
MHCII on IECs and hematopoietic granzymes regulate SI epithelial turnover. (A) Schematics showing the experimental setup. For the analysis of IEC turnover, mice were i.p. injected with EdU 72 hours before the analysis. (B-C) Analysis of the epithelial turnover in SFB+ MHCII^ΔIEC^ and cre-negative littermate mice. (B) Flow cytometry analysis of EdU+ IECs. FACS plots on the left show representative gating and numbers beside gates show frequency from all IECs. Plot on the right shows z-score batch-normalized frequency of EdU+ IECs. n=5-6. (C) Microscopic analysis of EdU+ cells. Representative staining images are shown on the left (DAPI-blue, EdU-green). Plot on the right shows statistical overview of EdU+ IECs/villus. 20-24 villi from 5-6 mice were analyzed. (D-E) Analysis of the epithelial turnover in BM chimeric animals. WT littermate recipients received either WT or Gzma^-/-^ Gzmb^-/-^ BM. After 6 weeks, mice were used for the analysis of SI epithelium turnover. (D) Flow cytometry analysis of EdU+ IECs. FACS plots on the left show representative gating and numbers beside gates show frequency from all IECs. Plot on the right shows z-score batch-normalized frequency of EdU+ IECs. n=5. (E) Microscopic analysis of EdU+ cells. Representative staining images are shown on the left (DAPI-blue, EdU-green). Plot on the right shows statistical overview of EdU+ IECs/villus. 24-39 villi from 3-5 mice were analyzed. Scale bar in (C) and (E) represents 100 µm. Horizontal lines in (B-E) show mean±SEM. Data in (B-E) was tested by Student’s t-test. *, p<0.05; ***, p<0.001; ****, p<0.0001.

We have shown that SFB induce strong expression of granzymes in SI-LP and IEL T-cell populations and that MHCII on the intestinal epithelium regulates the numbers of Gzmb^+^ SFB-specific iIEL. To test if granzymes can directly affect the epithelial turnover in SFB colonized animals, we next generated BM-chimeric animals, where WT SFB^+^ recipients were transplanted with either wild-type or Gzma^-/-^ and Gzmb^-/-^ (double knockout) BM. Six weeks after the BM transplantation, we analyzed the epithelial turnover in chimeric mice, using EdU pulse in a similar fashion as in Figure 7A. Importantly, animals lacking the hematopoietic expression of granzymes phenotypically matched the animals with epithelial deletion of MHCII (Figures 7D-E).

Collectively, this set of data suggests that SFB increase the turnover of the intestinal epithelium through the induction of granzyme producing iIELs that act in a MHCII-navigated manner.

## Discussion

Our study describes a previously uncharacterized type of immune response against SFB, a commensal bacterium of *Clostridia* taxon living in close association with IECs. This immune response is strictly age-dependent and starts by SFB-driven induction of IFNγ-expression in intestinal T_H_1 cells in a cDC1-dependent manner. Subsequently, the T_H_1-derived IFNγ induces massive expression of MHCII on IECs. In the ileum, which is the main site of SFB growth, MHCII expression is not restricted only to immature IECs (stem cells and TA), but also extends to mature enterocytes of higher villus zones. In parallel, SFB colonization induces differentiation of SFB-specific CD4^+^ T-cell clones into granzyme-expressing iIELs, which progressively accumulate in the intestine. Finally, we show that SFB-specific iIEL response ultimately regulates the turnover of IECs in MHCII and IEL’
ss granzyme-dependent manner. While previous studies described the importance of T_H_17 response against SFB (Ivanov *et al*., 2009; Ivanov et al., 2008), this study shows that in addition to this well-established mechanism, the immune reaction targeted against SFB also involves T_H_1 and cytotoxic iIELs responses. Spatially, the T_H_1 and T_H_17 responses against SFB are preferentially localized in the lamina propria, where IFNγ induces MHCII expression on IECs and IL-17 plus IL-22 cytokines induce recruitment of neutrophils (Flannigan *et al*., 2017), promote antimicrobial peptide production (Shih *et al*., 2014) and enhance epithelial healing (Pickert et al., 2009). In contrast, the cytotoxic iIEL response is localized in the intraepithelial space and propagates the renewal of the SFB-stressed gut epithelium by increasing its’ turnover rate.

While several recent studies described that MHCII presence in the intestinal epithelium is dependent on IFNγ (Al Nabhani *et al*., 2019; Koyama *et al*., 2019; Thelemann *et al*., 2014; Van Der Kraak *et al*., 2021), the source of this cytokine and the regulation of its production remained unclear. We have shown that under homeostatic conditions T_H_1 cells are the major source of IFNγ, responsible for the induction of MHCII on IECs. Our results stress that while the expression of MHCII in the intestinal epithelium is both age-dependent and IFNγ-induced, IFNγ produced during weaning is not sufficient for the long-lasting MHCII expression on IECs. Our study suggests that aside from T_H_17, SFB also induce a potent IFNγ -producing T_H_1 response, which is kept in equilibrium with the T_H_17 response through a feedback loop. This notion is supported by the fact that while IFNγ is the main inducer of MHCII expression in IECs, IL-17 acts in an opposite manner. Moreover, IFNγ production by T_H_1 cells is completely dependent on the cDC1 population. In line with our observation, cDC1 were shown to control a T_H_1 response to the recently discovered strain of *Cryptosporidium* that behaves as a commensal in WT mice (Russler-Germain *et al*., 2021), suggesting that mechanisms described in our study may be widely used in response to intestinal microbes whose survival depends on tight association with IECs (Atarashi *et al*., 2015; Chen and LaRusso, 2000)

The tight association with the host’
ss IECs might even be the defining factor necessary for the induction of an immune response described here, that is based on MHCII expression on IECs coupled with induction of iIELs. Indeed, it was shown previously that while infection by another IEC-associated bacterium - the *Citrobacter rodentium* (Takahashi et al., 2021), is well tolerated in WT mice, a large fraction of their MHCII^ΔIEC^ counterparts would not survive in the course of two weeks post infection (Jamwal *et al*., 2020). Interestingly, *Citrobacter rodentium* belongs to the group of attaching and effacing bacteria and often serves as the animal model for human enterotoxigenic *E. coli* infections (Collins et al., 2014; Wiles et al., 2006). This further highlights the importance of MHCII expression on the IECs in regulation of immune response against bacteria living tightly associated with IECs.

In the broader scope of MHCII’s role on IECs, there have been some seemingly conflicting reports regarding the role of MHCII in the maintenance of intestinal homeostasis in mice. Specifically, while MHCII expression on IECs protects the host from infectious colitis (Jamwal *et al*., 2020), it was found to aggravate chemical and T-cell induced colitis (Jamwal *et al*., 2020; Tuganbaev *et al*., 2020). Similarly, while MHCII on IECs was found to protect the host from the development of intestinal cancer (Beyaz *et al*., 2021), its expression on IECs was suggested to serve as the target for allogenic T-cells and thus drive graft-versus-host disease (Koyama *et al*., 2019). We believe that the mechanistic insight we show in this study can clarify this issue.

We have shown that one of the major functions of MHCII on IECs under homeostatic conditions is the regulation of numbers of iIEL specific for microbiota antigens in the intraepithelial space of the small intestine. Importantly, we were able to show by lineage tracing experiments that iIELs are generated from SFB-specific CD4^+^ conventional T-cells. SFB-specific iIELs start to express molecules necessary for tight association with the IECs and granzymes and thus bear cytotoxic potential. Moreover, SFB-specific iIELs are transcriptionally undistinguishable from *bona-fide* IELs. Thus, our study highlights the plasticity of the CD4^+^ T-cell clones, which can switch from mere helpers and coordinators to true fighters bearing cytotoxic potential. Furthermore, both MHCII on the IECs and hematopoietic expression of granzymes regulate the turnover of the intestinal epithelium.

It is thus possible to hypothesize that the immune system evolved a specific strategy, that is responsible for controlling bacterial strains tightly associated with IECs. The control is based on MHCII navigation of the cytotoxic, bacterium-specific iIEL response which is able to increase the IECs turnover. This hypothesis is in line with reported observations since such a mechanism would be beneficial if the immune system fights invading pathogens (Jamwal *et al*., 2020) or keeps in check commensal to prevent its overgrowth (Russler-Germain *et al*., 2021). On the other hand, it could cause severe damage, when the immune system is targeting gut epithelial cells without discrimination (in a state of graft-versus-host disease) (Koyama *et al*., 2019), or if the cytotoxic response to commensals gets out of control (like in inflammatory bowel disease) (Jamwal *et al*., 2020; Tuganbaev *et al*., 2020). It can also explain why the absence of MHCII on IECs can predispose them to malignant transformation (Beyaz *et al*., 2021) since the abrogation of immune surveillance by IEL and the disruption of possible cytotoxic elimination of (pre)malignant cells can be precisely the causative factor.

Mechanism we describe here may also have an outreach to antiviral intestinal responses. While a recent study showed that SFB protects the host from rotavirus infection by increasing epithelial turnover and shedding of infected cells (Shi et al., 2019), in another recent report authors show that iIEL derived from CD4^+^ T-cells protect the host from the infection with intestinal viral infections in IFNγ-dependent manner (Parsa et al., 2022). Our data suggest that protective effect of SFB can act thought the IFNγ-dependent induction of MHCII on IECs and concomitant switch of conventional SFB-specific CD4^+^ T-cells to iIEL with cytotoxic potential that can convey the protection from the virus. Thus, the immune response described here, and its defective regulation may have devastating (patho)physiologic consequences ranging from impaired clearance of invasive epithelia-associated bacteria to chronic intestinal inflammation or intestinal cancer.

## Materials and methods

### Mice

All mice used in this study were on C57Bl/6J background. Mice were used irrespectively of their sex since we observed not sex-depended differences in the analysis we performed. Three types of genetically identical C57B/6J WT mice were used. Two colonies of these C57B/6J wild-type mice were housed locally but in two separate barriers, since those two showed phenotypical differences, they were analyzed separately (see Results section). We also used C57Bl/6J animals shipped directly from Jackson Laboratories (JAX: 000664). SFB negative status of these animals was confirmed by qPCR and Gramm-staining from their faces. To keep the microbiota of these animals identical to that present in JAX, mice were directly transfer to the Trexler-type plastic isolator in Laboratory of Gnotobiology straight after shipping and maintained isolated further on. These mice were provided with autoclaved tap water, and rodent diet (Ssniff V1126-000) and bedding were sterilized by irradiation 50 kGy (Bioster, Veverská Bitýška, Czech Republic). All other mouse strains were housed under standard SPF conditions in the individually ventilated cages. R26-fl-STOP-fl-DTA (B6.129P2-Gt(ROSA)26Sortm1(DTA)Lky/J, 009669) (Voehringer *et al*., 2008), H2-Ab1^fl/fl^ (MHCII-fl, B6.129×1-H2-Ab1tm1Koni/J, 013181) (Hashimoto et al., 2002), Rag1^-/-^ (B6.129S7-Rag1tm1Mom/J, 002216) (Mombaerts et al., 1992), CD90.1 (B6.PL-Thy1a/CyJ, 000406) and SFBtg (Tg(Tcra,Tcrb)2Litt/J, 027230) (Yang *et al*., 2014) mouse strains were purchased from JAX. Xcr1-Cre (Xcr1-IRES-iCre-GSG-2A-mTFP) (Wohn *et al*., 2020) animals were kindly provided by Dr. Bernard Malissen. Vil1-CreERT (B6.Cg-Tg(Vil1-cre/ERT2)23Syr/J) (el Marjou et al., 2004) mice were kindly provided by Dr. Vladimir Korinek. Gzma^-/-^ Gzmb^-/-^ (B6.Cg-Gzmatm1Simn Gzmbtm1Ley) (Simon et al., 1997) mice were kindly provided by Julián Pardo, University of Zaragoza, Spain. All mice were given standard rodent high energy breeding diet and reverse osmosis-filtered water *ad libitum*. Mice were kept at 12 h/12 h light/dark cycle; temperature and relative humidity were maintained at 22 ± 1°C and 55 ± 5%, respectively. All experiments were approved by the and the ethical committee of the Faculty of Science of the Charles University, the ethical committee of the Institute of Molecular Genetics of the Czech Academy of Science and the ethical committee of the Institute of Microbiology of the Czech Academy of Science.

### Cell isolations

For the isolation of SI epithelium, SI-LP and SI-IEL cells we euthanized the mouse by cervical dislocation, dissected complete SI and maintained it in PBS on ice. SI was then cleaned from mesenteric fat and luminal contents were rinsed with ice-cold PBS using syringe. Equal portions (6 cm) of each of small intestinal segments – duodenum, jejunum and ileum were then cut and the rest of the tissue was discarded. PP were excised carefully and the remaining tissue was cut open longitudally and separated to 2 cm long pieces. This tissue served as the input material for the downstream isolation methods.

For the isolation of SI epithelium, tissue was incubated in HBSS without Ca^2+^ and Mg^2+^ supplemented with 3% fetal bovine serum (Gibco, FBS) and 2mM EDTA (stripping buffer) on ice for 90 minutes, followed by 10 seconds of vortexing. Tissue was transferred to a fresh stripping buffer and incubation/vortexing was repeated several times, with 30-minutes incubation time and the stripping buffer containing pieces of the epithelium were collected. For the analysis of complete SI-epithelium (crypts and villi) second and third fractions were pooled and used further. For the analysis of crypt-resident epithelial cells, fourth and fifth fractions were passed through 100µm nylon filter and pooled together. Epithelium was then collected by centrifugation (3 min, 300g, 4 ºC) and digested by TrypLE express (Gibco) for 1 minute at 37ºC and vortexed. Cell suspension was passed through 100µm filter, washed with stripping buffer and used for downstream applications.

For the isolation of SI-IEL cells SI tissue was incubated in the stripping buffer at 37ºC for 20 minutes. This procedure was repeated one more time and the buffer containing the epithelium was pooled, collected by centrifugation, digested by TrypLE express for 1 minute at 37ºC and vortexed. Cell suspension was passed through 100µm filter and washed with stripping buffer. IELs were enriched by Percoll (Cytiva) gradient centrifugation. Specifically, cells were resuspended in 4ml of 40% Percoll in RPMI medium (Gibco) supplemented with 3% FBS. This suspension was then underlaid with 2ml of 80% Percoll in RPMI medium supplemented with 3% FBS. This gradient was centrifuged for 20 minutes at 800g at 21ºC, without brake. IEL cells were collected from the interface, washed with stripping buffer and used for downstream applications.

For the isolation of SI-LP cells SI tissue was stripped of the epithelium in the same way as for the isolation of SI-IEL. The remaining tissue was then cut to ∼2mm pieces and incubated in enzymatic cocktail at 37ºC for 1h. This cocktail contained Colleagenase D (Roche, 1 mg/ml) and DNAse I (Roche, 40 U/ml) dissolved in RPMI medium supplemented with 3% FBS. Tissue was then vortexed for 10 seconds and passed through 100µm filter. The remaining tissue was meshed on 100µm cell strainer. Cells collected by mashing were pooled with the flow-through, washed with the stripping buffer and collected by centrifugation. LP cells were isolated by Percoll gradient centrifugation. Cells were resuspended in 4ml of 40% Percoll in RPMI medium (Gibco) supplemented with 3% FBS. This suspension was then underlaid with 2ml of 80% Percoll in RPMI medium supplemented with 3% FBS. This gradient was centrifuged for 20 minutes at 800g at 21ºC, without brake. Cells were collected from the interface, washed with stripping buffer and used for downstream applications.

For the isolation of T-cells for the adoptive transfer spleen, axillary, brachial, inguinal, and cervical lymph nodes were dissected. These tissues were cut to ∼2mm pieces and incubated for 1h at 37ºC in the enzymatic cocktail containing Colleagenase D (Roche, 1 mg/ml) and DNAse I (Roche, 40 U/ml) dissolved in RPMI medium supplemented with 3% FBS. Tissue was then vortexed for 10 seconds and passed through 100µm filter. The remaining tissue was meshed on 100µm cell strainer. Cells collected by mashing were pooled with the flow-through, washed with the stripping buffer and collected by centrifugation. T-cells were isolated by Naive CD4+ T Cell Isolation Kit (Miltenyi Biotec) in accordance with the manufacturer’
ss instructions.

For the analysis of PP and mLN, these tissues were dissected and incubated for 1h at 37ºC in the enzymatic cocktail containing Colleagenase D (Roche, 1 mg/ml) and DNAse I (Roche, 40 U/ml) dissolved in RPMI medium supplemented with 3% FBS. Tissue was then vortexed for 10 seconds and passed through 100µm filter. The remaining tissue was meshed on 100µm cell strainer. Cells collected by mashing were pooled with the flow-through, washed with the stripping buffer and collected by centrifugation.

For the isolation of bone marrow, donor mice were euthanized by cervical dislocation and their femurs and tibias were dissected. Joints were cut-out from the bone and bone marrow was isolated by ice-cold PBS rinse. Bone marrow was collected by centrifugation, single cell suspension was generated by pipetting and passed through 100µm filter.

### Mice treatment

For the SFB colonization, freshly collected feces of the SFB mono-associated mice were collected and frozen immediately in 50% glycerol in PBS+0.05% Cystein hydrochloride. Presence of SFB bacteria and absence of contamination was assessed by Gramm-staining of the fecal smear and qPCR analysis with SFB-specific primers. Acceptor mice were fed with 50 µl of thawed SFB suspension at 21 and 22 days of age. These mice were analyzed two weeks later. For the IFNγ blockade, mice were i.p. injected with 250 µg of anti-IFNγ antibody (XMG1.2, Biolegend) or the same amount of isotype control in 100 µl of PBS. For injection and analysis timepoints please see respective figure legends. For the NK/NKT cell depletion, mice were i.p. injected with 25 µg of Nk1.1 antibody (PK136, BioXcell) at 21, 23, 25, 29, 31 and 33 days of age and analyzed at 35 days of age. For the inducible MHCII depletion on IECs, Vil1-CreER x MHCII-fl/fl mice were i.p. injected with 1 mg of tamoxifen (Sigma) in 50 µl of sunflower oil at 21 and 22 days of age. For the adoptive transfer of T-cells, mice were intravenously (i.v.) injected with 5×10^6^ of purified CD4^+^ T-cells in 100 µl of sterile PBS. These mice were analyzed 2 weeks later. For the generation of BM chimeras, the recipient mice were irradiated (6 Gy, X-RAD 225XL; Accela) and i.v. injected with 10^7^ donor BM cells in 100 µl of PBS. To ensure better survival of recipients, they were maintained on gentamycin (1 mg/ml) in drinking water for a week post irradiation. After antibiotics treatment, recipient mice were fed with 50 µl of thawed SFB suspension 1 and 2 days after the termination of antibiotics treatment. These mice were analyzed 4-6 weeks after the irradiation. For the analysis of epithelial turnover, mice were i.p. injected with EdU (1.25 mg/mouse) in 100 µl of PBS 72 hours before the analysis.

### Flow cytometry

Cells were incubated in the antibody cocktail in the stripping buffer (HBSS without Ca^2+^ and Mg^2+^ supplemented with 3% FBS and 2mM EDTA) for 20 minutes on ice, washed and either directly analyzed by flow cytometry or processed further for intracellular staining. For the staining of intracellular epitopes, cells were fixed using Foxp3 / Transcription Factor Staining Buffer Set (Invitrogen) for 30 minutes and stained with the cocktail of antibodies at room temperature in the Foxp3 staining kit permeabilization buffer for 30 minutes. For the staining of cytokines and granzymes, cells were incubated in the stimulation cocktail (phorbol myristate acetate (20 ng/ml, Sigma), ionomycin (1 μg/mL, Sigma) and Brefeldin A (10 μg/mL, Invitrogen) in RPMI supplemented with 3% FBS) for 3-4 hours at 37ºC prior to the antibody staining. For the discrimination of live cells, Hoechst33258 or Fixable Viability Dye eFluo 506 (eBioscience) was used. For the staining of EdU+ IECs, Click-iT EdU Flow Cytometry Assay Kit AF488 (Life Technologies) was used in accordance with the manufacturer’s instructions. Samples were analyzed using LSRII or Symphony FACS flow cytometers (both BD Biosciences). Flow cytometry-based cell sorting was done using FACSAria IIu cell sorter (BD Biosciences). Flow cytometry data were analyzed using FlowJO software (v10.8.1, BD Biosciences).

### Fluorescence microscopy

Small intestine and colon tissues were dissected, and their luminal contents were washed with ice-cold PBS by syringe. Tissues were then fixed in 4% formaldehyde for 2 hours at room temperature. Cryoprotection was performed by the incubation in 30% sucrose in PBS over-night at 4ºC. Tissues were submerged in O.C.T. medium (Agar Scientific) and snap-frozen on dry ice. Frozen tissues were stored at -80°C and cut to 12µm sections using CM1950 cryostat (Leica). Slides for immunofluorescence microscopy were incubated in 4% PFA for 10 minutes, washed 3 times in PBS, permeabilized in 100% methanol at -20°C for 10 minutes, washed 3 times in PBS again, incubated in blocking buffer (PBS, 2.5% FBS, 2.5% BSA) for 1 hour at room temperature and stained with primary antibody over-night at 4°C. Following the overnight stain, the slides were washed 3 times with PBS, incubated with secondary antibody for 2 hours at room temperature, washed 3 times with PBS, mounted with Vectashield with DAPI (Vector Laboratories), and imaged using Dragonfly spinning-disk fluorescence microscope (Andor) or widefield fluorescent microscope DMI8 (Leica). For the EdU staining on microscopic sections, frozen sections were allowed to come to room temperature, washed three times with PBS and blocked with 5% BSA in PBT. After that, the manufacturers protocol was followed using Click-iT EdU Flow Cytometry Assay Kit A488 (Life Technologies).

### qPCR analysis

qPCR was used for the routine screening of SFB presence. For this analysis bacterial DNA was isolated either from feces or from 1 cm of mouse terminal ileum using QIAmp Fast DNA Stool kit (QIAGEN). qPCR was performed using SYBR Green I qPCR Master mix on LC480II Light cycler (Both Roche). Following primers were used for this analysis: SFB (F: GACGCTGAGGCATGAGAGCAT, R: GACGGCACGGATTGTTATTCA), universal bacterial probe (F: ACTCCTACGGGAGGCAGCAGT, R: ATTACCGCGGCTGCTGGC)(Yin et al., 2013). Water sample instead of bacterial was used as a negative control. Relative abundance of SFB was calculated using the relative quantification method(Pfaffl, 2001). In SFB monoisolates as well as in SFB monocolonized mice SFB abundance was always equal to 1. In SFB positive mice with complex microbiota SFB abundance ranged from ∼10^−5^ to 10^−2^. In SFB-free mice the reaction with SFB primers formed no valid PCR product.

### Single cell RNA sequencing

For the droplet-based scRNA-seq, freshly isolated cells were FACS-sorted using CD45, EpCAM and Hoechst33258 staining. LP and IEL cells were sorted as live CD45^+^ cells, whereas epithelial cells were sorted as EpCAM^+^ CD45^-^ live cells. SI-IECs and SI-LP from WT 3-, 4-, 5- and 6-weeks old mice were isolated and sorted separately but SI-IECs and SI-LP from each timepoint were pooled before the 10x sequencing pipeline. Similarly, SI-IECs and SI-IEL from JAX and SFB colonized JAX mice were isolated and sorted separately but cells from each mice were pooled before the 10x sequencing pipeline. Separation of IECs from LP/IEL cells was performed bioinformatically based on the combination of UMAP clustering and canonic hematopoietic (*Ptprc*) and epithelial (*Epcam*) marker genes. SI-LP cells from JAX and SFB colonized JAX mice cells were processed separately.

The viability of cells was assessed by trypan blue and counted in an automated TC20 cell counter (Bio-Rad) prior libraries preparation. Single-cell RNA-seq libraries were prepared using Chromium controller instrument and Chromium Next Gem single-cell 3’ reagent kit version 3.1 (both 10X Genomics) according to the manufacturer’s protocol. The quality and quantity of the resulting cDNA and libraries were determined using Agilent 2100 Bioanalyzer (Agilent Technologies). The libraries were sequenced using NextSeq 500 instrument (Illumina) in accordance with the manufacturer’s protocol with mRNA fragment read length of 54 bases. We used 10X Genomics Cell Ranger software suite (version 4.0.0) to quantify gene-level expression based on GRCm38 assembly (Ensembl annotation version 98) (Cunningham et al., 2019).

Well-based scRNA-seq was done using SORT-Seq platform (Muraro et al., 2016). Freshly isolated SI-LP and SI-IEL cells (as described above) were stained with antibodies recognizing TCRβ, CD4, CD8α, CD8β, CD90.1, CD90.2 and Vβ14 subunit of TCR. SFBtg T-cells were sorted as live TCRβ+ CD90.1+ Vβ14+ cells. Host cells were sorted as live TCRβ+ cells. All the markers were recorded while sorting and later integrated with scRNA-seq data using indexes generated during cell sorting. Cells were sorted into 384-well plates (Single cell discoveries) and snap frozen on dry ice straight after sorting. The CEL-Seq2 protocol was used for library preparation by Single cell discoveries (Muraro *et al*., 2016). The libraries were sequenced using NextSeq 500 instrument (Illumina) according to the manufacturer’s protocol.

### 16S Sequencing

For the analysis of the ileum microbiome, 1 cm of tissue was dissected, cut open longitudinally and immediately frozen at -80°C. DNA was then extracted using the QIAamp PowerFecal DNA Kit (QIAGEN).

For the amplification reaction of bacterial 16S rDNA, primers covering V4-V5 region were used (forward: GGATTAGATACCCTGGTAGT, reverse: CACGACACGAGCTGACG) (Fliegerova et al., 2014). The thermal cycling conditions included an initial denaturation of 10 min at 95 °C followed by 25 cycles of 30s at 95 °C, 30s at 57 °C, and 30s at 72 °C. Sequencing libraries were prepared using NEBNext® Fast DNA Library Prep Set for Ion Torrent™ (New England BioLabs), diluted to the same concentration and pooled. The sequencing template was prepared in a One Touch 2 instrument while the sequencing itself took place in an Ion Torrent PGM platform.

Raw sequences were processed using the Qiime2 2020.6. Qiime2 embedded DADA2 was used for checking quality, trimming, dereplication, denoising and removal of chimeras. Similarly, a VSEARCH-based consensus classifier was used to assign taxonomy against the Greengenes database, version 13_8 with 97 % identity. Sequences were rarefied at a minimum sequencing depth.

### Bioinformatic analysis

All the bioinformatic analyses were done using R (version 4.1.1). For the analysis of droplet-based scRNA-seq datasets, 10x pipeline was used to generate read-count matrices. This data served as an input for all the downstream analysis. First, the ambient RNA was removed using SoupX R package (v 1.5.2). Cells that contained more than 5% of mitochondrial RNA from their total mapped reads were removed. This step was followed by the separation of IECs from LP (3-, 4-, 5- and 6-weeks old WT mice) or IEL (JAX and SFB colonized mice) cells which were sequenced together. For this UMAP clustering and the expression of canonic hematopoietic (*Ptprc*) and epithelial (*Epcam*) marker genes, was used to. Thus, cells of hematopoietic or epithelial origin were defined, separated and processed separately in all downstream analyses. Further data processing was done using standard Seurat R package (v 4.0.4) pipeline. Briefly, cells were clustered, and their cell types annotated based on canonic markers. When cluster contained a mixed population, this approach was repeated, and the cluster was further subdivided to populations representing unique cell types. Thus, we identified cell types, which served as the main components for further downstream analysis.

Similar to droplet-based scRNA-seq datasets, we processed plate-based scRNA-seq datasets by a standard Seurat R package (v 4.0.4) pipeline.

For the analysis of relative bacterial abundance read counts generated by 16S sequencing were analyzed using DESeq2 R package. Plain p-value and log2 fold change values were used for the analysis.

### Statistics

Aside from sequencing datasets, all the remaining data was tested using GraphPad Prism 9.

## Acknowledgements

Research in the Dobeš laboratory (JD, TB, KK, IP) is kindly supported by the Czech Science Foundation JUNIOR STAR grant (No. 21-22435M), Czech Science Foundation grant (No. 22-30879S) and by Charles University PRIMUS grant (No. Primus/21/MED/003). MS and DŠrůtková are supported by the Czech Science Foundation JUNIOR STAR grant (No. 21-19640M). DF, MD and JB are supported by Grant #20-30350S and #22-30879S from the Czech Science Foundation (GA CR). MK is supported by Operational Programme Research, Development and Education project (No. CZ.02.1.01/0.0/0.0/16_019/0000785).

We are thankful to Martina Krausová and Šárka Kocourková from the Genomics and Bioinformatics core facility of the Institute of Molecular Genetics in Prague, Czech Republic, for the preparation of single cell libraries and to Zdeněk Cimburek from the Flow Cytometry Core Facility of the Institute of Molecular Genetics in Prague, Czech Republic for suggestions related to cell sorting. We thank the Animal Facility in Krč personnel of the Institute of Molecular Genetics in Prague, Czech Republic for their excellent maintenance of our animal colonies and Šárka Maisnerová from the Laboratory of Gnotobiology, Institute of Microbiology in Nový Hrádek, Czech Republic for the germ-free and SFB negative mice maintenance. We are thankful to Bernard Malissen (Centre d’Immunologie de Marseille Luminy, Marseille, France) and Julián Pardo (University of Zaragoza, Zaragoza, Spain) for providing Xcr1-cre and granzyme double knockout animals.

## Author contributions

TB, DF, JA and JD initiated the project. TB and JD designed vast majority of experiments, critically discussed the data, and wrote and revised the manuscript in close collaboration. TB executed and analyzed majority of the experiments, JD executed some of the experiments and acquired the funding. JA and DF critically discussed the results, revised the manuscript and contributed with critically important reagents. MS critically contributed to the design of gnotobiotic experiments and together with DŠrůtková provided SFB monoisolates, performed all SFB colonization experiments and handled all gnotobiotic animals. KK performed majority of microscopy analysis and provided technical support for the project. MD performed some of the experiment, carried out majority of mouse genotyping and provided technical support for the project. DSchierová performed microbiome analysis. JB performed animal irradiation, critically discussed the data and the manuscript. IP provided support in some of the experiments. OB-N, YG and IS performed some of the experiments. MK provided technical expertise in scRNA-seq experiments.

## Data availability

Data will be made available in the final version.

## Figure legends

**Figure S1.Markers defining cell populations in scRNA-seq analysis of SI-LP from 3-, 4-, 5- and 6-weeks old WT mice, related to Figure 1**

(A) Scatter heatmap of markers defining first division of cell populations to B cells, plasma cells, myeloid cells and other lymphoid cells.

(B) Scatter heatmap of markers defining subpopulations of lymphoid cells.

(C) Scatter heatmap of markers defining subpopulations of myeloid cells.

**Figure S2.Scatter heatmap of markers defining cell populations in scRNA-seq analysis of SI-IEC from 3-, 4-, 5- and 6-weeks old WT mice, related to Figure 2**

**Figure S3.Age-dependent changes in EEC and SI stem cell compartments, related to Figure 2**

(A-C) Analysis of SI-EEC subpopulations in 3-, 4-, 5- and 6-weeks old WT mice.

(A) UMAP of EEC subclustering and cell type annotation defined in this dataset.

(B) Expression of canonic markers in EEC clusters defining EEC subtype annotations.

(C) Changes in Log2 FC of frequencies of EEC subpopulations through mouse age.

(D-G) Analysis of SI stem cell and TA cells subpopulations in 3-, 4-, 5- and 6-weeks old WT mice.

(D) UMAP of Stem/TA subclustering and cell type annotation defined in this dataset.

(E) Expression of canonic markers in Stem/TA clusters defining Stem cells subtype annotations.

(F) Expression of MHCII molecules in Stem/TA clusters through mouse age.

(G) Changes in Log2 FC of frequencies of Stem/TA clusters through mouse age.

**Figure S4.Age-dependent changes in enterocyte subpopulations based on region of origin and position on the villus, related to Figure 2**

(A) UMAP of enterocyte subclustering and annotation based on the region or origin.

(B) Violin plot of the expression of proximal (duodenum) and distal (ileum) markers defining the annotation of enterocytes subclusters.

(C) Expression of MHCI and MHCII and genes responsible for their antigen processing in enterocyte subclusters based on their region of origin.

(D-F) Subclustering of enterocytes from duodenum (D), jejunum (E) and ileum (F) and their annotation to zone on the villus.

(G-I) Expression of the markers defining zone on the villus in enterocyte subcluster from duodenu (G), jejunum (H) and ileum (I).

(J-L) Expression of MHCI and MHCII and genes responsible for their antigen processing in enterocyte subcluster from duodenum (J), jejunum (K) and ileum (L).

**Figure S5.Overview of antigen presentation-associated genes in IEC subpopulation at 6 weeks, related to Figure 2**

(A) Heatmap of MHCI and MHCII genes expression in individual IEC subpopulations from 6-weeks-old animals.

(B) Heatmap of proteasomes, cathepsins and transport proteins responsible for MHCI and MHCII antigen processing genes expression in individual IEC subpopulations from 6-weeks-old animals.

(C) Heatmap of both classical and non-classical costimulatory molecules expression in individual IEC subpopulations from 6-weeks-old animals.

**Figure S6.Kinetics of MHCII expression on IECs, related to Figure 2**

(A-B) Flow cytometry analysis of MHCII expression on IECs at 2-7 weeks of age.

(A) Representative FACS plots.

(B) Plot showing the overview of the frequency of MHCII+ IECs from 3 independent litters, where each litter is represented in each timepoint, n=3-5 per timepoint. Horizontal lines show mean±SEM.

(C) Immunofluorescence microcopy of SI from 3-(top row) or 5-weeks-old (bottom row) animals. EpCAM (white), MHCII (red), Lysozyme 1 (green) and CD45 (blue) were stained. Scale bar represents 50 μm.

**Figure S7.SFB induced changes in the epithelial compartment, related to Figure 3**

(A) Flow cytometry analysis of Paneth cell frequency from all IECs (CD45-EpCAM+). Plots show representative gating and the statistical overview of the data. n=6.

(B-D) scRNA-seq analysis of EEC compartment after SFB colonization of JAX mice.

(B) UMAP of EEC subclustering and cell type annotation defined in this dataset.

(C) Expression of canonic markers in EEC clusters defining EEC subtype annotations.

(D) Changes in Log2 FC of frequencies of EEC subpopulations after SFB colonization.

(E-H) scRNA-seq analysis of SI stem cell and TA cells subpopulations after SFB colonization of JAX mice.

(E) UMAP of Stem/TA subclustering and cell type annotation defined in this dataset.

(F) Expression of canonic markers in Stem/TA clusters defining Stem cells subtype annotations.

(G) Changes in Log2 FC of frequencies of Stem/TA clusters after SFB colonization.

(H) Expression of MHCII molecules in Stem/TA clusters after SFB colonization.

**Figure S8.SFB induced changes in the enterocyte subpopulations, related to Figure 3**

(A) UMAP of enterocyte subclustering and annotation based on the region or origin.

(B) Violin plot of the expression of proximal (duodenum) and distal (ileum) markers defining the annotation of enterocytes subclusters.

(C) Expression of MHCI and MHCII and genes responsible for their antigen processing in enterocyte subclusters based on their region of origin.

(D-F) Subclustering of enterocytes from duodenum (D), jejunum (E) and ileum (F) and their annotation to zone on the villus.

(G-I) Expression of the markers defining zone on the villus in enterocyte subcluster from duodenum (G), jejunum (H) and ileum (I).

(J-L) Expression of MHCI and MHCII and genes responsible for their antigen processing in enterocyte subcluster from duodenum (J), jejunum (K) and ileum (L).

**Figure S9.SFB induced T-cell responses, related to Figure 4**

(A) Representative gating strategy used for the discrimination of basic T-cell populations.

(B-C) Flow cytometry analysis of cytokine expression in intestinal T-cells in 5-weeks-old JAX mice, half of which was colonized with SFB at 21 and 22 days of age.

(B) Analysis of IFNγ expression in SI-LP CD8^+^ T-cells. Plot on the left shows frequency of IFNγ^+^ CD8^+^ T-cells. Plot on the right shows MFI of IFNγ in all CD8 T-cells. n=7.

(C) Flow cytometry analysis of cytokine expression in SI-IEL T-cells. Top row shows frequency IFNγ^+^, MFI of IFNγ, frequency of IL-17^+^ and the frequency of IFNγ^+^ IL-17^+^ in CD4 T-cells. Bottom row shows frequency of IFNγ^+^ in CD8^+^ T-cells. Plot on the right shows MFI of IFNγ in all CD8^+^ T-cells. n=7. Horizontal lines in (B-C) show mean±SEM. Data in (B-C) was tested by Student’s t-test and p values are shown. MFI results are shown as batch-normalized using z-score.

**Figure S10.Mechanisms regulating epithelial MHCII expression on the cellular level, related to Figure 4**

(A) Analysis of IFNγ producers. IFNγ expression is shown across all defined cell types in SI-LP at 6 weeks of age, when MHCII expression on IECs is high.

(B) Flow cytometry analysis of IFNγ producers in SI-LP of 6-weeks-old mice. Representative gating of IFNγ^+^ cells to basic T- and NK cell populations is shown. Numbers beside gates represent frequencies.

(C) Nk1.1 mediated depletion of SI-LP NK and NKT cells. Mice were treated with either isotype antibody or Nk1.1 depleting antibody at 21, 23, 25, 29, 31 and 33 days of age and analyzed at 35 days of age FACS in the top row show representative NK (Nk1.1+ TCRβ-) and NKT (Nk1.1+ TCRβ+) gating and their frequencies. Plots in the bottom row show the efficiency of the depletion. n=5

(D) Flow cytometry of MHCII expression on SI-IECs in Nk1.1 depleted and littermate control mice. FACS plots on the left show representative gating and frequency of MHCII+ IECs. Plot on the right shows statistic overview of the data. n=5. Horizontal lines in (C-D) show mean±SEM. Data in (C-D) was tested by Student’s t-test. *, p<0.05p values >0.05 are shown.

**Figure S11.cDC1 controlled T-cell responses, related to Figure 4**

(A) Efficiency of cDC1 depletion in Xcr1-Cre x R26-fl-STOP-fl-DTA (XCR1-DTA) mice. Plots on the left show representative gating of all XCR1+ cells from mLN in XCR1-DTA and littermate control (Xcr1-Cre^-^). Numbers beside gates show counts. Plot on the right shows statistical overview of cDC1 counts. n=4-7.

(B) T-cell responses in the intestine of XCR1-DTA mice. Top row shows counts of CD4^+^ and CD8^+^ T-cells in SI-LP of XCR1-DTA and littermate control mice. Bottom row shows the frequency of IFNγ^+^ cells and IFNγ MFI in CD8+ SI-LP T-cells. n=4-7. Horizontal lines in (A-B) show mean±SEM. Data in (A-B) was tested by Student’s t-test. ****, p<0.0001; p values >0.05 are shown.

**Figure S12.Markers defining cell populations in LP and IEL in JAX and SFB colonized JAX mice, related to Figure 5**

(A) Canonic markers defining the cell type annotation of clusters in SI-LP from JAX and SFB colonized JAX mice.

(B) Canonic markers defining the cell type annotation of clusters in SI-IEL from JAX and SFB colonized JAX mice.

**Figure S13.SFB mediated induction of cytotoxic phenotype in LP and IEL T-cell subpopulations, related to Figure 5**

(A) Representative gating of CD4, CD8αα and CD8αβ T-cell populations. Numbers beside gates represent frequencies.

(B-E) Flow cytometry analysis of T-cell counts, frequency of Gzmb+ cells and z-score batch-normalized MFI of Gzmb in SI-LP CD8αβ T-cells (B), SI-LP CD8αα T-cells (C), SI-IEL CD8αβ T-cells (D) and SI-IEL CD8αα T-cells (E) from JAX and SFB colonized JAX mice. n=7. Data was batch-normalized using z-score. Horizontal lines in (B-E) show mean±SEM. Data in (B-E) was tested by was tested by Student’s t-test. **, p<0.01; p values >0.05 are shown.

**Figure S14.Cell type definition in well-based scRNA-seq of intestinal T-cells, related to Figure 6**

(A) Representative gating of CD90.1+ T-cells carrying TCR specific for SFB-specific antigen (SFBtg). Numbers beside gates represent frequencies.

(B) UMAP of well-based scRNA-seq analysis of pooled SI-LP and SI-IEL T-cells from WT and MHCII^ΔIEC^ mice, which were injected with naïve CD4+ SFBtg T-cells 2 weeks prior to analysis.

(C) Expression of canonic markers defining cell type annotation of populations shown in (B).

(D) Scatter heatmap of RNA (top row) and protein (bottom row) expression of CD4, CD8α and CD8β in well-based scRNA-seq analysis of SI-IEL and SI-LP T-cells.

**Figure S15.T-cell responses controlled by MHCII on the intestinal epithelium, related to Figure 6**

(A-D) Congenically marked SFBtg BM was mixed with feeder WT BM in 5:95 ratio and i.v. injected to irradiated WT or MHCII^ΔIEC^ hosts. MHCII^ΔIEC^ hosts were induced with tamoxifen at 21 and 22 days of age and the BM transplantation was carried out at 4 weeks of age of the acceptors. These mice were analyzed 4-6 weeks after BM transplantation.

(A) Flow cytometry analysis of SFBtg T-cell from SI-LP.

(B) Flow cytometry analysis of WT SI-LP T-cells divided according to their CD4, CD8αα or CD8αβ identity.

(C) Flow cytometry analysis of SFBtg T-cell from SI-IEL.

(D) Flow cytometry analysis of WT SI-IEL T-cells divided according to their CD4, CD8αα or CD8αβ identity. Data was batch-normalized using z-score. Horizontal lines show mean±SEM. n=4-6. Data was tested by Student’s t-test. *, p<0.05; **, p<0.01; p values >0.05 are shown.

