## supplemental figures for "Epithelial antigen presentation controls commensal-specific intraepithelial T-cells in the gut"

### Figure S1

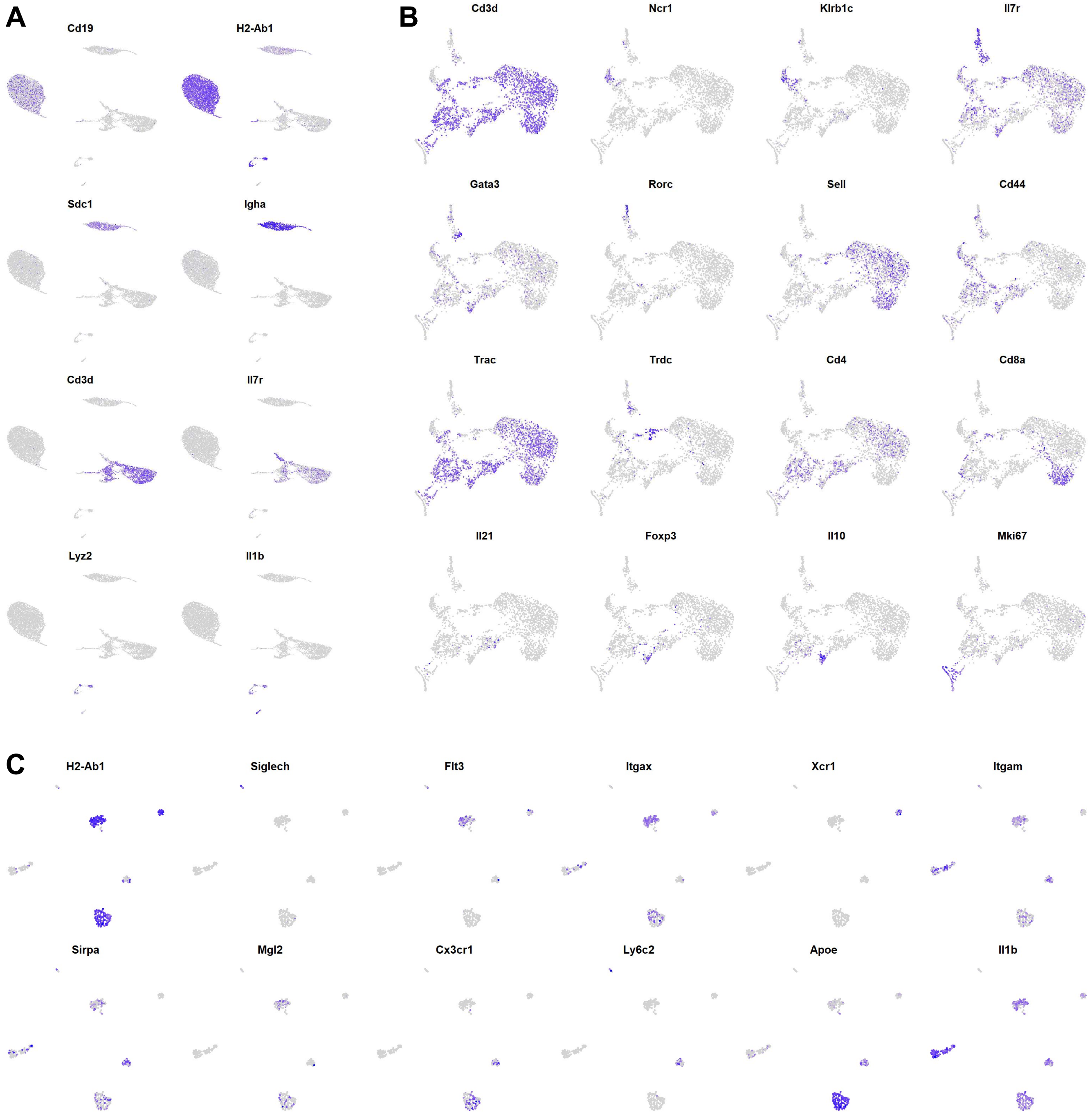
A
Cd19
H2-Ab1
Sdc1
Igha
Cd3d
Il7r
Lyz2
Il1b
B
Cd3d
Ncr1
Klrb1c
Il7r
Gata3
Rorc
Sell
Cd44
Trac
Trdc
Cd4
Cd8a
Il21
Foxp3
Il10
Mki67
C
H2-Ab1
Siglech
Flt3
Itgax
Xcr1
Itgam
Sirpa
Mgl2
Cx3cr1
Ly6c2
Apoe
Il1b

#### Figure S2

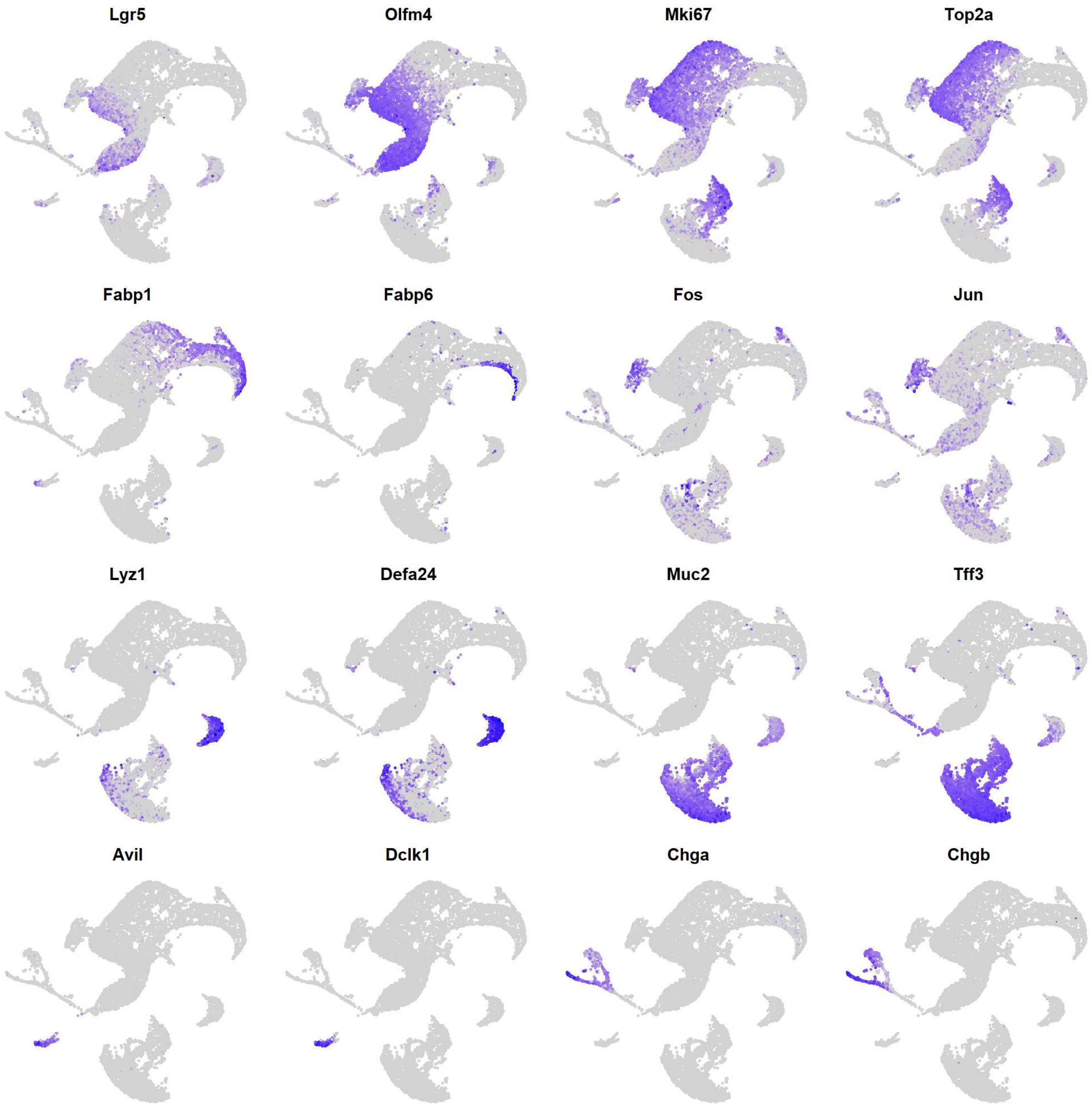
Lgr5
Olfm4
Mki67
Top2a
Fabp1
Fabp6
Fos
Jun
Lyz1
Defa24
Muc2
Tff3
Avil
Dclk1
Chga
Chgb

#### Figure S3

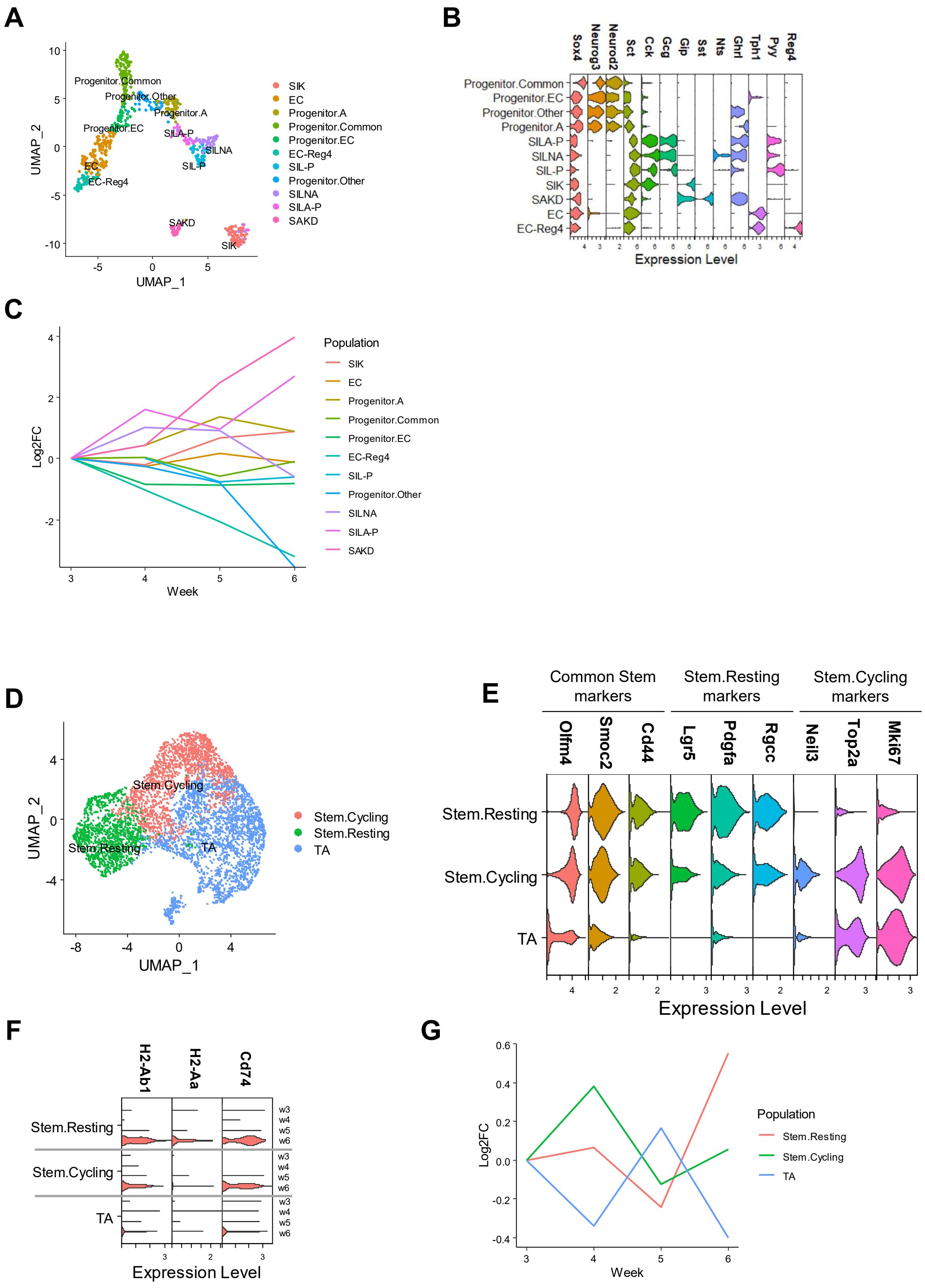

A
B
C
D
E
F
G
UMAP_1
UMAP_2
Progenitor.Common
Progenitor.Other
Progenitor.A
Progenitor.EC
SILA-P
SILNA
SIL-P
EC
EC-Reg4
SAKD
SIK
Expression Level
Population
Log2FC
Week
Stem.Cycling
Stem.Resting
TA
Common Stem markers
Stem.Resting markers
Stem.Cycling markers
Olfm4
Smoc2
Cd44
Lgr5
Pdgfa
Rgcc
Neil3
Top2a
Mki67
H2-Ab1
H2-Aa
Cd74
Sox4
Neurog3
Neurod2
Sct
Cck
Gcg
Gip
Sst
Nts
Ghrl
Tph1
Pyy
Reg4

### Figure S4

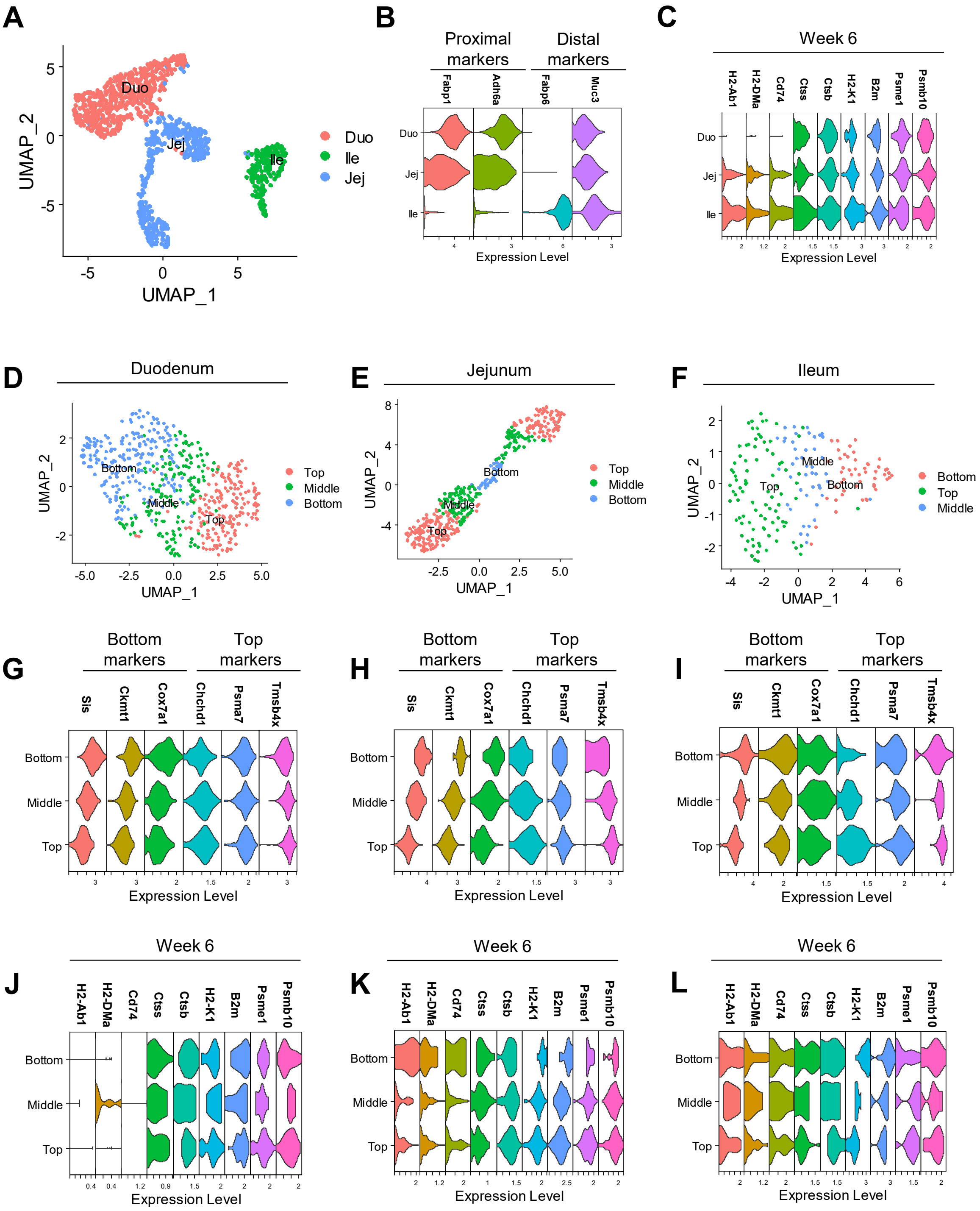

A
B
C
D
E
F
G
H
I
J
K
L
UMAP_1
UMAP_2
Duo
Ile
Jej
Proximal markers
Distal markers
Fabp1
Adh6a
Fabp6
Muc3
Week 6
H2-Ab1
H2-DMa
Cd74
Ctss
Ctsb
H2-K1
B2m
Psme1
Psmb10
Expression Level
Duodenum
Jejunum
Ileum
Top
Middle
Bottom
Bottom markers
Top markers
Sis
Ckmt1
Cox7a1
Chchd1
Psma7
Tmsb4x

### Figure S5

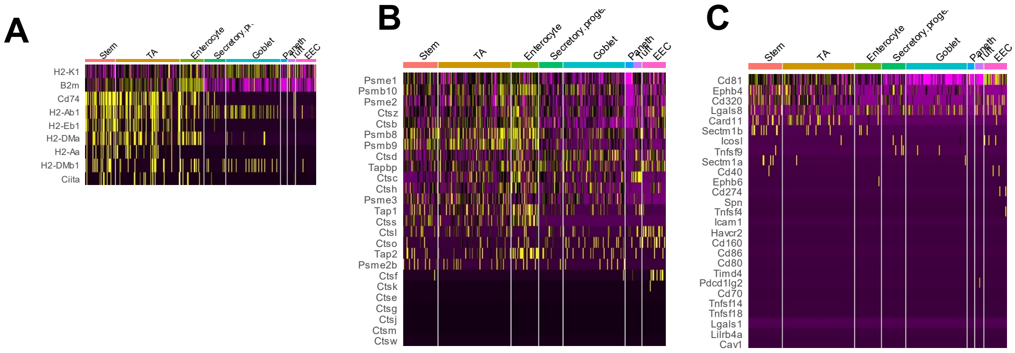

A
Stem
TA
Enterocyte
Secretory p..
Goblet
Paneth
Tuft
EEC
H2-K1
B2m
Cd74
H2-Ab1
H2-Eb1
H2-DMa
H2-Aa
H2-DMb1
Ciita
B
Stem
TA
Enterocyte
Secretory proge
Goblet
Paneth
Tuft
EEC
Psme1
Psmb10
Psme2
Ctsz
Ctsb
Psmb8
Psmb9
Ctsd
Tapbp
Ctsc
Ctsh
Psme3
Tap1
Ctss
Ctsl
Ctso
Tap2
Psme2b
Ctsf
Ctsk
Ctse
Ctsg
Ctsj
Ctsm
Ctsw
C
Stem
TA
Enterocyte
Secretory proge
Goblet
Paneth
Tuft
EEC
Cd81
Ephb4
Cd320
Lgals8
Card11
Sectm1b
Icosl
Tnfsf9
Sectm1a
Cd40
Ephb6
Cd274
Spn
Tnfsf4
Icam1
Havcr2
Cd160
Cd86
Cd80
Timd4
Pdcd1lg2
Cd70
Tnfsf14
Tnfsf18
Lgals1
Lilrb4a
Cav1

#### Figure S6

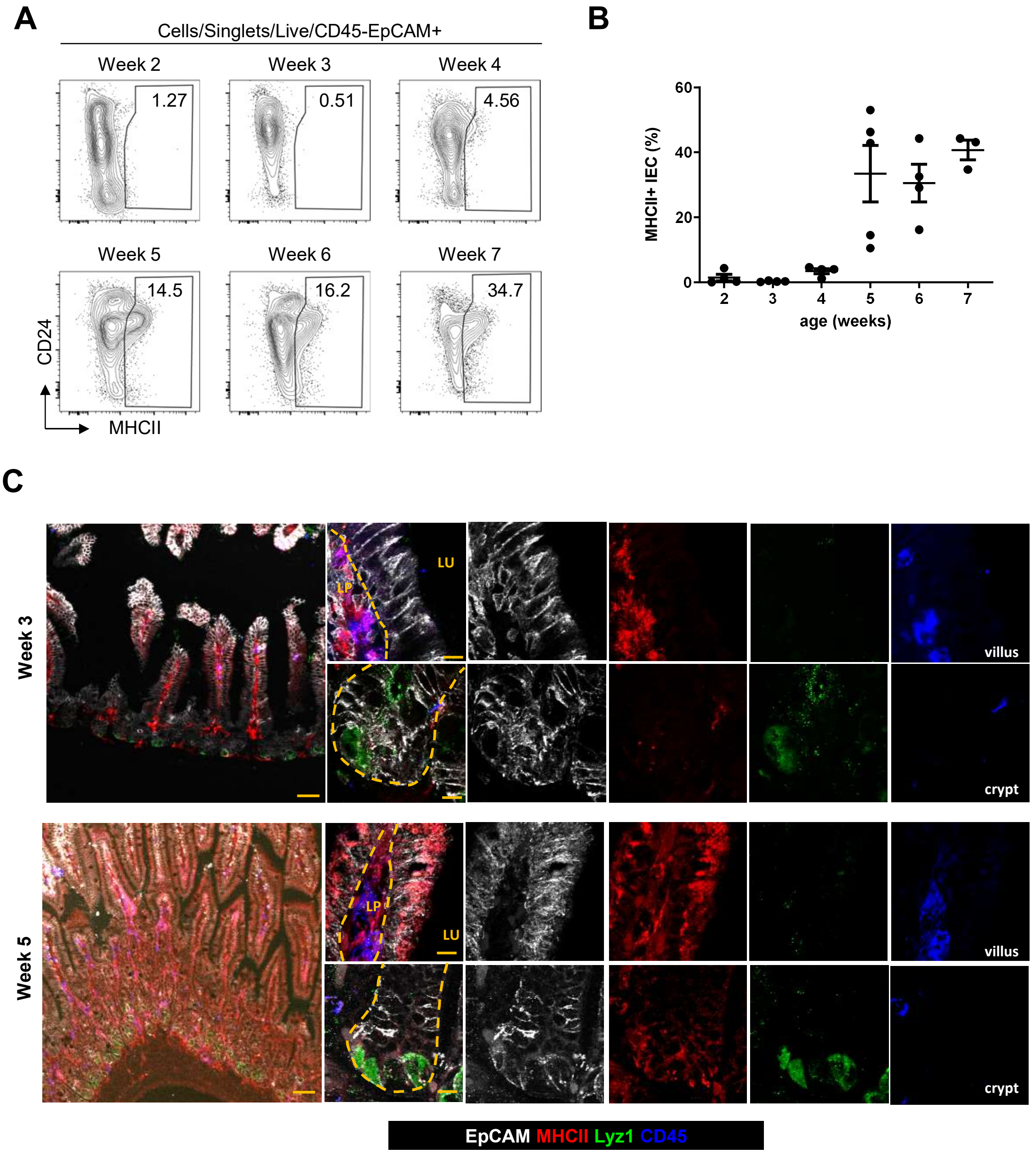
A
Cells/Singlets/Live/CD45-EpCAM+
Week 2
1.27
Week 3
0.51
Week 4
4.56
Week 5
14.5
Week 6
16.2
Week 7
34.7
CD24
MHCII
B
MHCII+ IEC (%)
60
40
20
0
2
3
4
5
6
7
age (weeks)
C
Week 3
LU
LP
villus
crypt
Week 5
LP
LU
villus
crypt
EpCAM MHCII Lyz1 CD45

### Figure S7

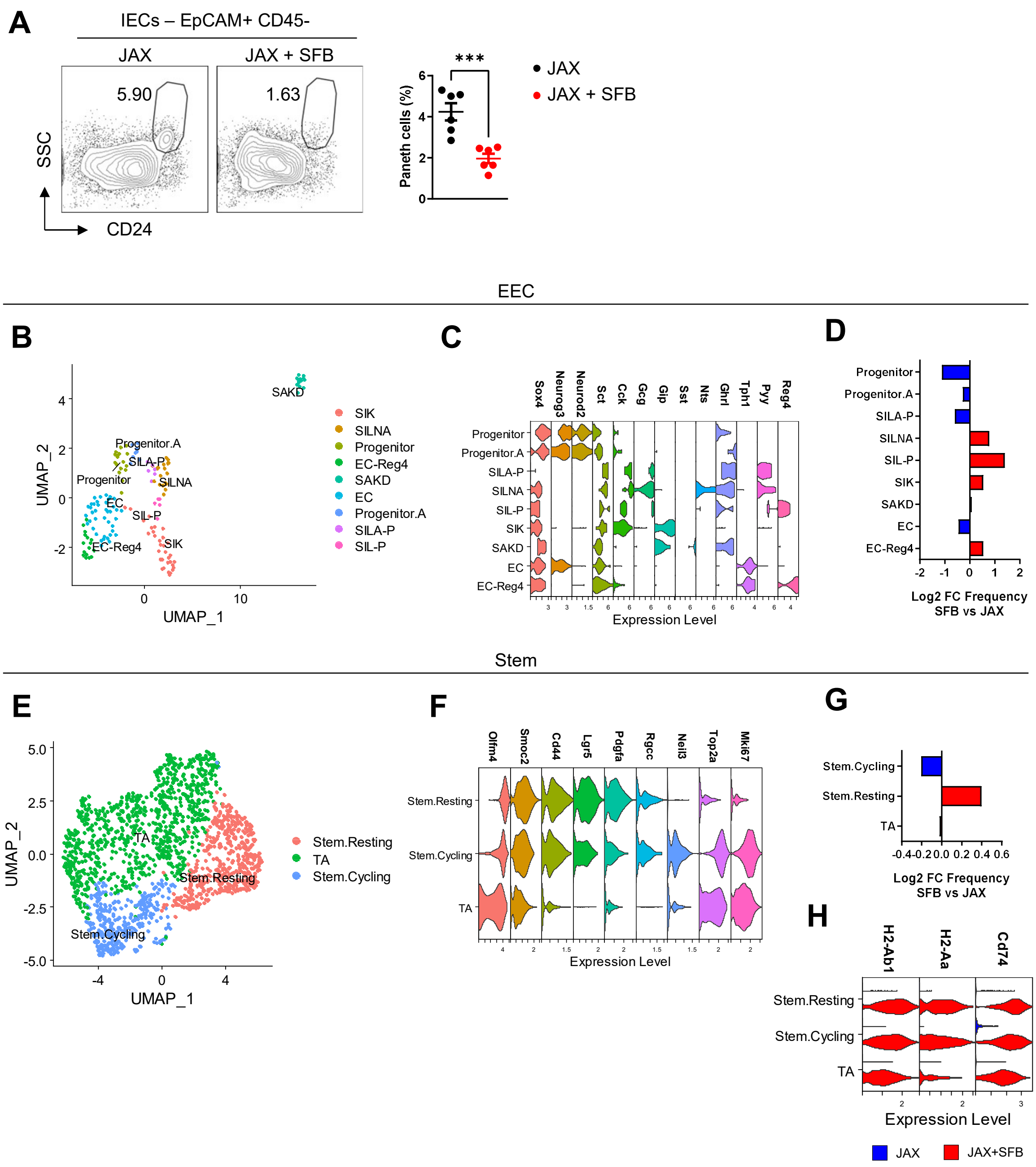
A
IECs – EpCAM+ CD45-
JAX
JAX + SFB
5.90
1.63
SSC
CD24
Paneth cells (%)
***
JAX
JAX + SFB
EEC
B
C
D
SIK
SILNA
Progenitor
EC-Reg4
SAKD
EC
Progenitor.A
SILA-P
SIL-P
UMAP_1
UMAP_2
Sox4
Neurog3
Neurod2
Sct
Cck
Gcg
Gip
Sst
Nts
Ghrl
Tph1
Pyy
Reg4
Expression Level
Log2 FC Frequency
SFB vs JAX
Stem
E
F
G
H
Stem.Resting
TA
Stem.Cycling
Olfm4
Smoc2
Cd44
Lgr5
Pdgfa
Rgcc
Neil3
Top2a
Mki67
H2-Ab1
H2-Aa
Cd74
JAX
JAX+SFB

### Figure S8

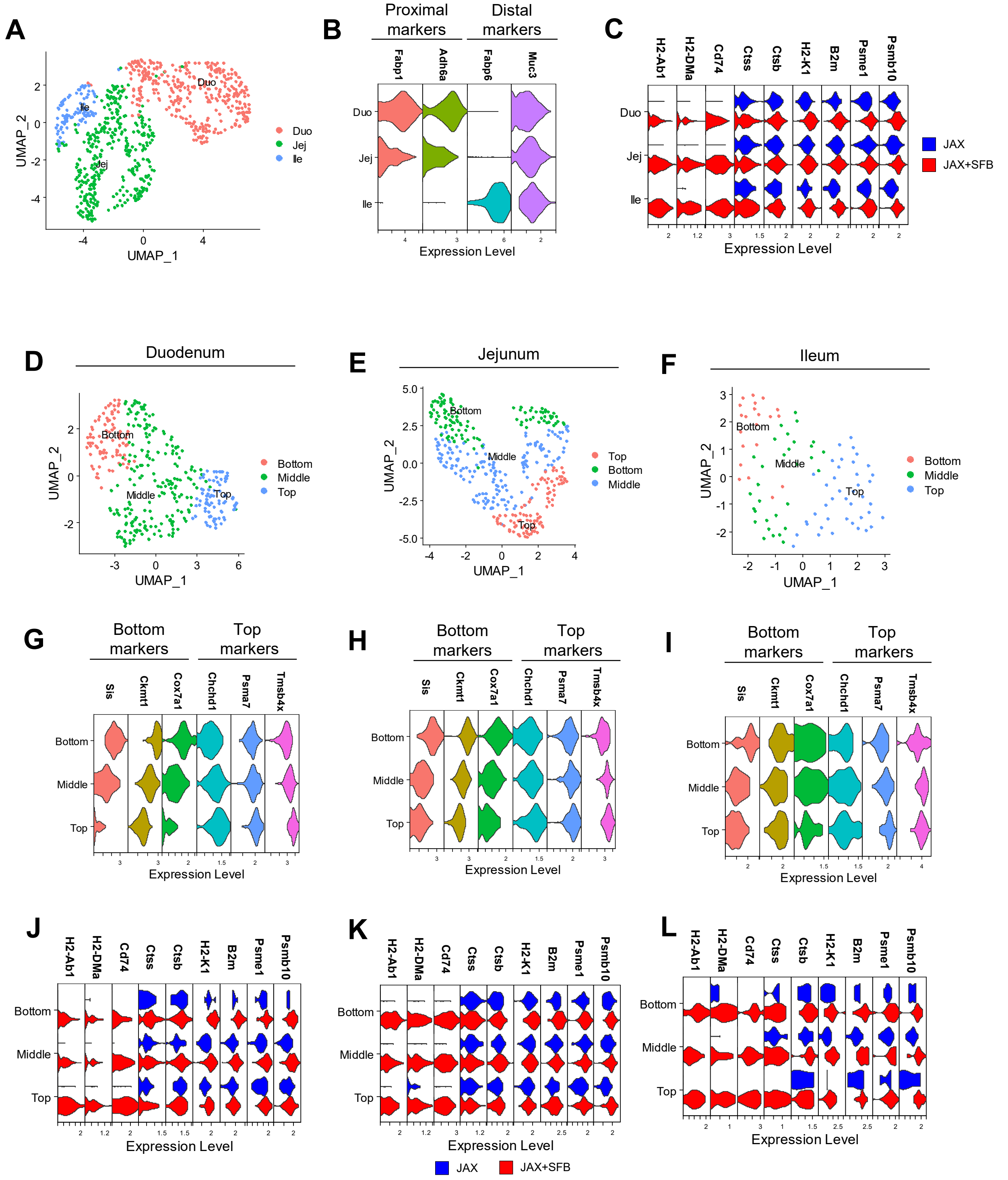

A
UMAP_1
UMAP_2
Duo
Jej
Ile
B
Proximal markers
Distal markers
Fabp1
Adh6a
Fabp6
Muc3
Expression Level
C
H2-Ab1
H2-DMa
Cd74
Ctss
Ctsb
H2-K1
B2m
Psme1
Psmb10
JAX
JAX+SFB
D
Duodenum
E
Jejunum
F
Ileum
Bottom
Middle
Top
G
H
I
Bottom markers
Top markers
Sis
Ckmt1
Cox7a1
Chchd1
Psma7
Tmsb4x
J
K
L

**Figure S9**

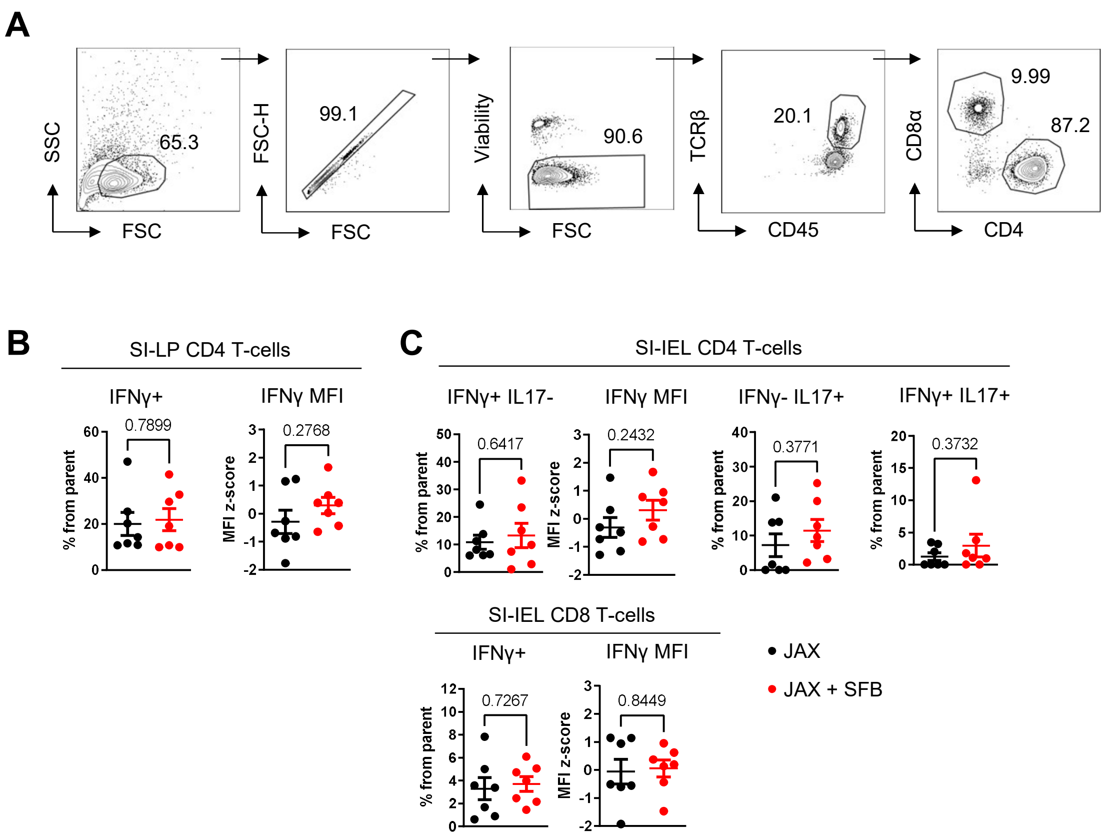
A
SSC
FSC
65.3
FSC-H
FSC
99.1
Viability
FSC
90.6
TCRβ
CD45
20.1
CD8α
CD4
9.99
87.2
B
SI-LP CD4 T-cells
IFNγ+
0.7899
% from parent
IFNγ MFI
0.2768
MFI z-score
C
SI-IEL CD4 T-cells
IFNγ+ IL17-
0.6417
IFNγ MFI
0.2432
IFNγ- IL17+
0.3771
IFNγ+ IL17+
0.3732
SI-IEL CD8 T-cells
IFNγ+
0.7267
IFNγ MFI
0.8449
JAX
JAX + SFB

**Figure S10**

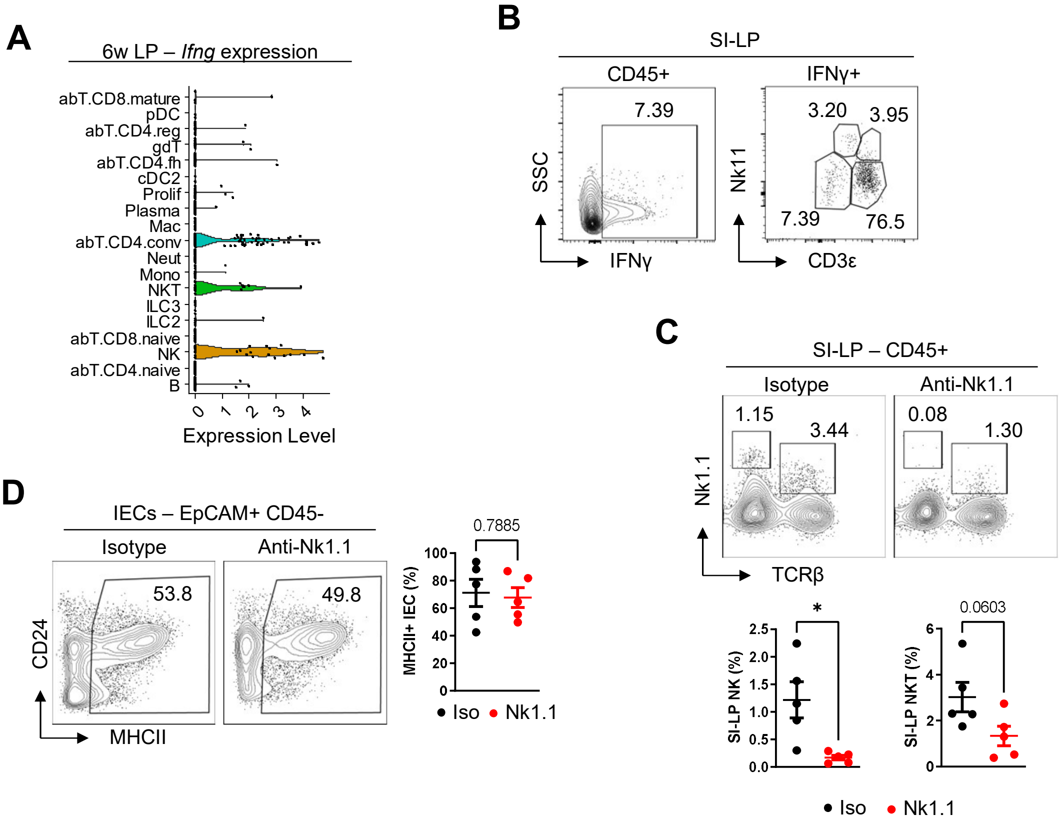
A
6w LP – Ifng expression
abT.CD8.mature
pDC
abT.CD4.reg
gdT
abT.CD4.fh
cDC2
Prolif
Plasma
Mac
abT.CD4.conv
Neut
Mono
NKT
ILC3
ILC2
abT.CD8.naive
NK
abT.CD4.naive
B
0
1
2
3
4
Expression Level
B
SI-LP
CD45+
IFNγ+
7.39
3.20
3.95
7.39
76.5
SSC
IFNγ
Nk11
CD3ε
C
SI-LP – CD45+
Isotype
Anti-Nk1.1
1.15
3.44
0.08
1.30
Nk1.1
TCRβ
*
0.0603
SI-LP NK (%)
SI-LP NKT (%)
● Iso
● Nk1.1
D
IECs – EpCAM+ CD45-
Isotype
Anti-Nk1.1
53.8
49.8
CD24
MHCII
0.7885
MHCII+ IEC (%)
● Iso
● Nk1.1

**Figure S11**

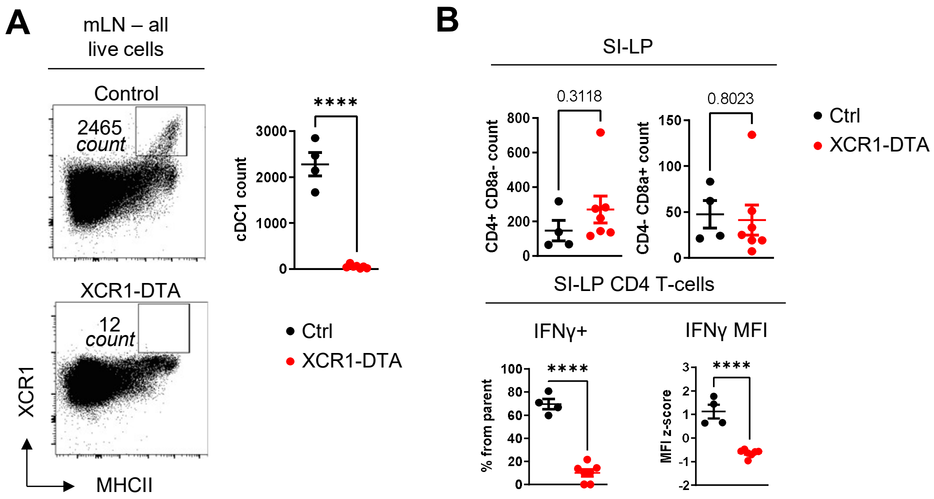
A
mLN – all live cells
Control
2465 count
XCR1-DTA
12 count
XCR1
MHCII
cDC1 count
****
3000
2000
1000
0
• Ctrl
• XCR1-DTA
B
SI-LP
0.3118
CD4+ CD8a- count
800
600
400
200
0
0.8023
CD4- CD8a+ count
150
100
50
0
• Ctrl
• XCR1-DTA
SI-LP CD4 T-cells
IFNγ+
% from parent
****
100
80
60
40
20
0
IFNγ MFI
MFI z-score
****
3
2
1
0
-1
-2

### Figure S12

A

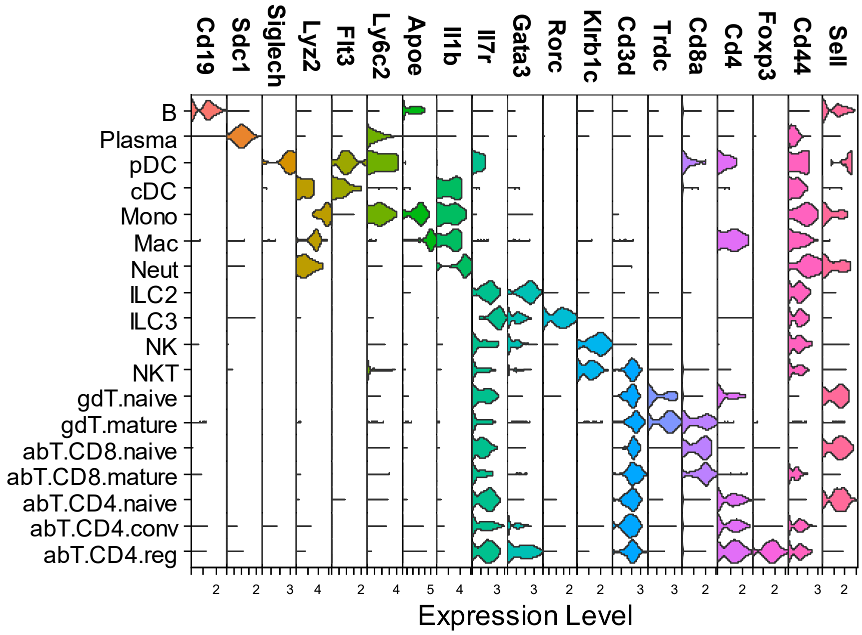


B

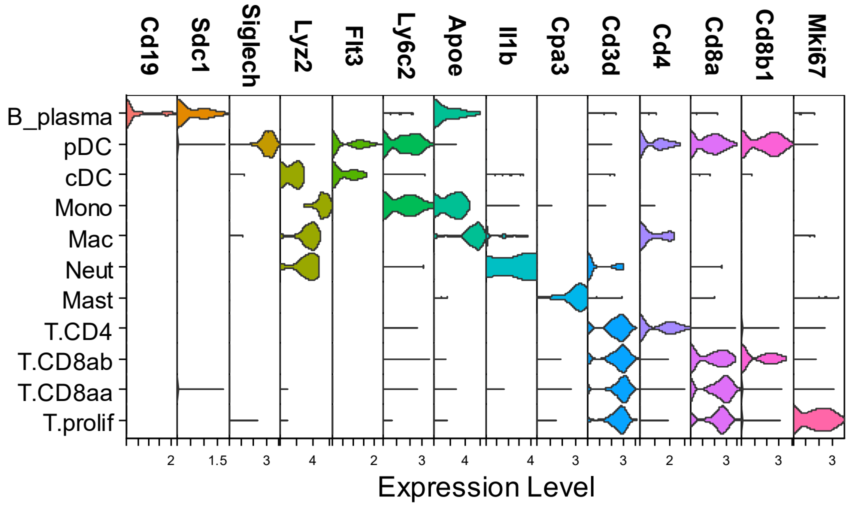

#### Figure S13

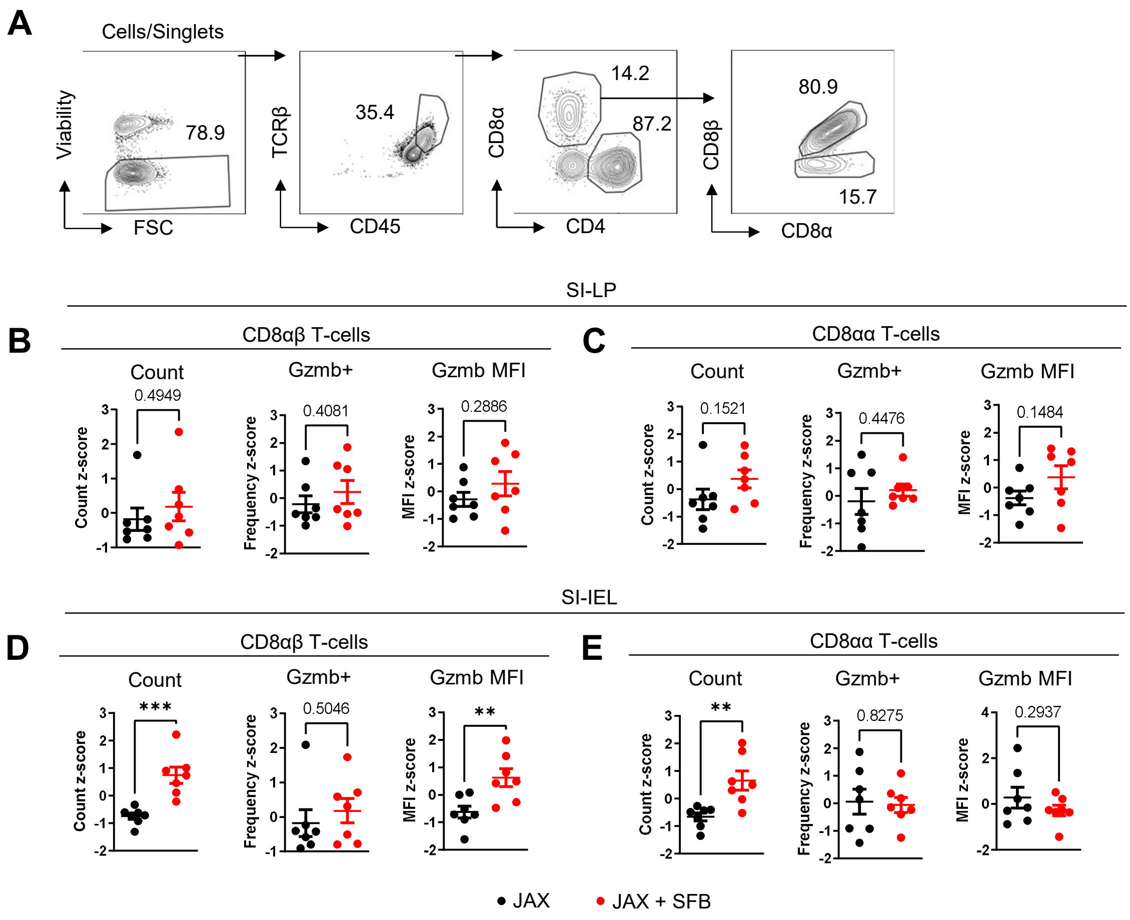
A
Cells/Singlets
Viability
FSC
78.9
TCRβ
CD45
35.4
CD8α
CD4
14.2
87.2
CD8β
CD8α
80.9
15.7
SI-LP
B
CD8αβ T-cells
Count
0.4949
Count z-score
Gzmb+
0.4081
Frequency z-score
Gzmb MFI
0.2886
MFI z-score
C
CD8αα T-cells
Count
0.1521
Count z-score
Gzmb+
0.4476
Frequency z-score
Gzmb MFI
0.1484
MFI z-score
SI-IEL
D
CD8αβ T-cells
Count
***
Count z-score
Gzmb+
0.5046
Frequency z-score
Gzmb MFI
**
MFI z-score
E
CD8αα T-cells
Count
**
Count z-score
Gzmb+
0.8275
Frequency z-score
Gzmb MFI
0.2937
MFI z-score
JAX
JAX + SFB

**Figure S14**

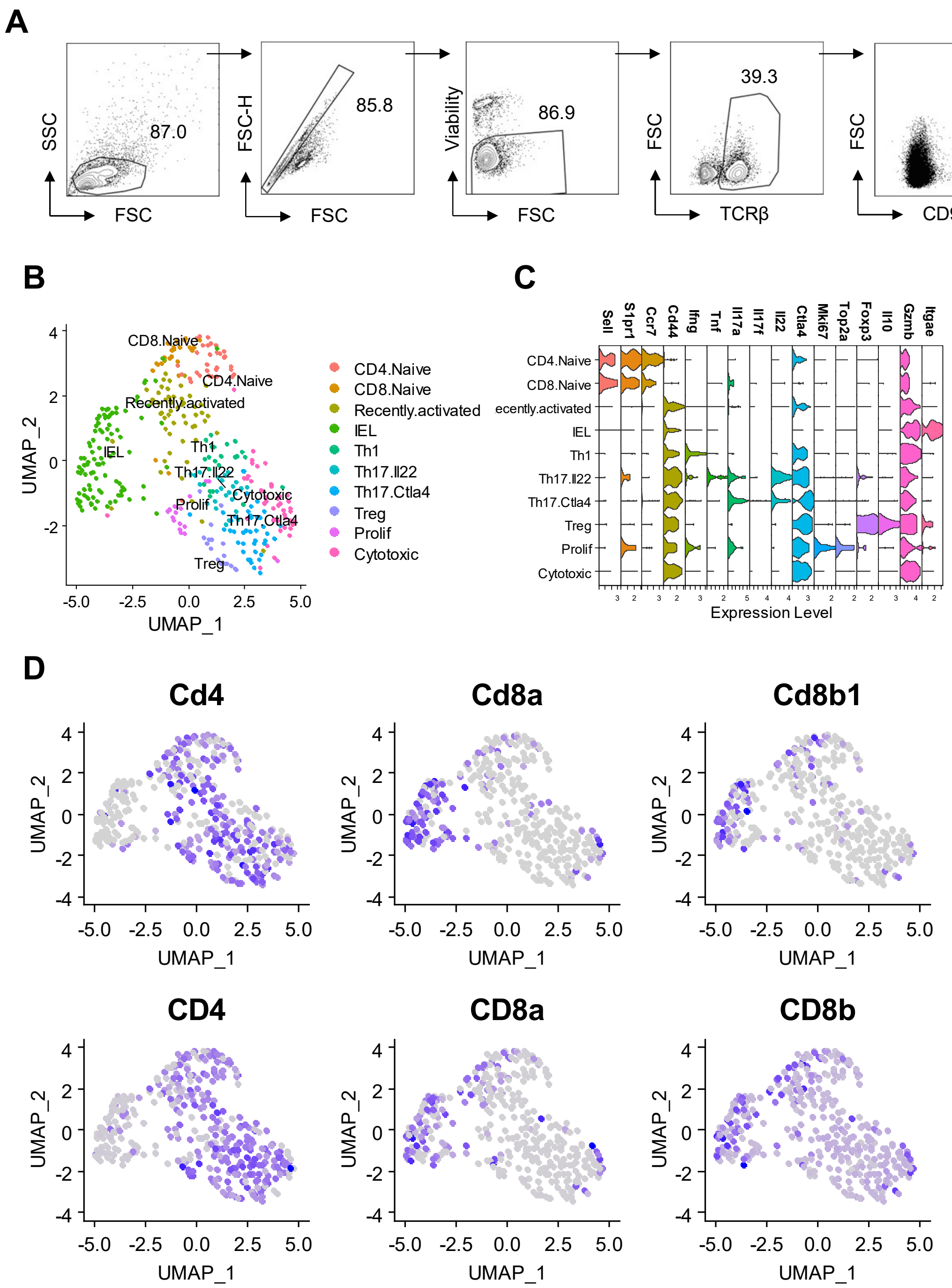
A
87.0
85.8
86.9
39.3
1.20
SSC
FSC
FSC-H
Viability
TCRβ
CD90.1
B
CD4.Naive
CD8.Naive
Recently.activated
IEL
Th1
Th17.Il22
Th17.Ctla4
Treg
Prolif
Cytotoxic
UMAP_1
UMAP_2
C
Sell
S1pr1
Ccr7
Cd44
Ifng
Tnf
Il17a
Il17f
Il22
Ctla4
Mki67
Top2a
Foxp3
Il10
Gzmb
Itgae
Expression Level
D
Cd4
Cd8a
Cd8b1
CD4
CD8a
CD8b

#### Figure S15

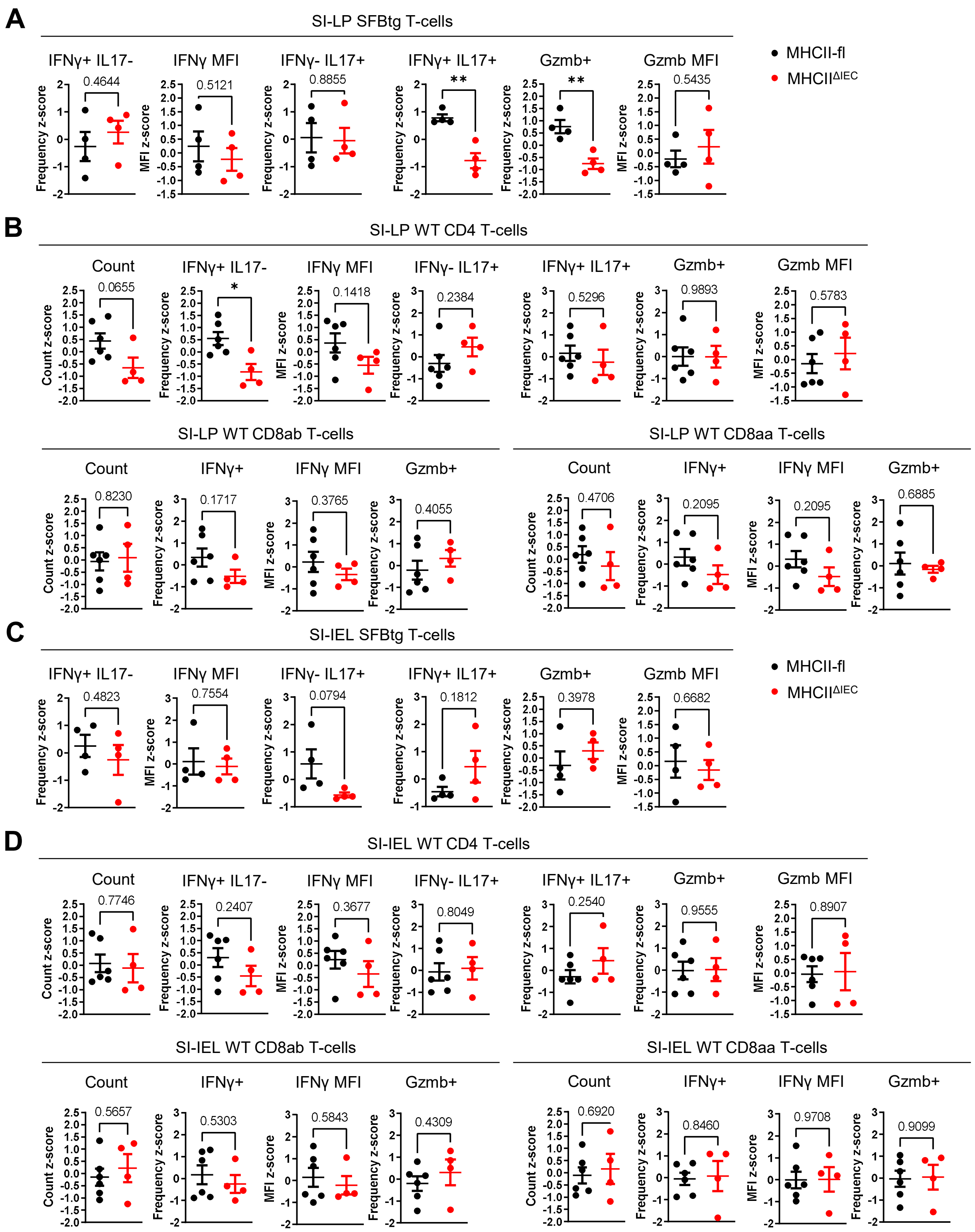
A
SI-LP SFBtg T-cells
IFNγ+ IL17-
IFNγ MFI
IFNγ- IL17+
IFNγ+ IL17+
Gzmb+
Gzmb MFI
0.4644
0.5121
0.8855
**
**
0.5435
MHCII-fl
MHCIIΔIEC
B
SI-LP WT CD4 T-cells
Count
IFNγ+ IL17-
IFNγ MFI
IFNγ- IL17+
IFNγ+ IL17+
Gzmb+
Gzmb MFI
0.0655
*
0.1418
0.2384
0.5296
0.9893
0.5783
SI-LP WT CD8ab T-cells
Count
IFNγ+
IFNγ MFI
Gzmb+
0.8230
0.1717
0.3765
0.4055
SI-LP WT CD8aa T-cells
Count
IFNγ+
IFNγ MFI
Gzmb+
0.4706
0.2095
0.2095
0.6885
C
SI-IEL SFBtg T-cells
IFNγ+ IL17-
IFNγ MFI
IFNγ- IL17+
IFNγ+ IL17+
Gzmb+
Gzmb MFI
0.4823
0.7554
0.0794
0.1812
0.3978
0.6682
MHCII-fl
MHCIIΔIEC
D
SI-IEL WT CD4 T-cells
Count
IFNγ+ IL17-
IFNγ MFI
IFNγ- IL17+
IFNγ+ IL17+
Gzmb+
Gzmb MFI
0.7746
0.2407
0.3677
0.8049
0.2540
0.9555
0.8907
SI-IEL WT CD8ab T-cells
Count
IFNγ+
IFNγ MFI
Gzmb+
0.5657
0.5303
0.5843
0.4309
SI-IEL WT CD8aa T-cells
Count
IFNγ+
IFNγ MFI
Gzmb+
0.6920
0.8460
0.9708
0.9099
Frequency z-score
MFI z-score
Count z-score
